# An Aurora B-RPA signaling axis secures chromosome segregation fidelity

**DOI:** 10.1101/2022.09.26.509563

**Authors:** Poonam Roshan, Sahiti Kuppa, Jenna R. Mattice, Vikas Kaushik, Rahul Chadda, Nilisha Pokhrel, Brunda Tumala, Brian Bothner, Edwin Antony, Sofia Origanti

## Abstract

Errors in chromosome segregation underlie genomic instability associated with cancers. Resolution of replication and recombination intermediates and protection of vulnerable single-stranded DNA (ssDNA) intermediates during mitotic progression requires the ssDNA binding protein Replication Protein A (RPA). However, the mechanisms that regulate RPA specifically during unperturbed mitotic progression are poorly resolved. RPA is a heterotrimer composed of RPA70, RPA32 and RPA14 subunits and is predominantly regulated through hyperphosphorylation of RPA32 in response to DNA damage. Here, we have uncovered a mitosis-specific regulation of RPA by Aurora B kinase. Aurora B phosphorylates Ser-384 in the DNA binding domain B of the large RPA70 subunit and highlights a mode of regulation distinct from RPA32. Disruption of phosphorylation markedly affects cell viability and causes defects in chromosome segregation with hypersensitivity to DNA damaging agents. Phosphorylation at Ser-384 remodels the protein interaction domains of RPA and facilitates formation of higher density filaments. Furthermore, phosphorylation impairs RPA binding to DSS1 that likely suppresses homologous recombination during mitosis by preventing recruitment of DSS1-BRCA2 to exposed ssDNA. We showcase a critical RPA-Aurora B signaling axis in mitosis that is essential for maintaining genomic integrity.

## Introduction

Maintaining genomic integrity during chromosome replication, condensation, and segregation relies on regulatory mechanisms that are distinct to each phase of the cell cycle.^1,2^ Protection of transiently exposed ssDNA throughout the cell cycle is achieved through binding of Replication Protein A (RPA).^3^ RPA also serves as a protein-interaction hub to recruit other proteins onto DNA and coordinates almost all DNA metabolic processes including replication, repair, recombination, and telomere maintenance.^4-6^ RPA performs several essential functions in the cell. It binds to ssDNA with high affinity (K_D_ <10^−10^ M) and protects it from degradation by exo- and endonucleases.^3^ Formation of RPA-ssDNA complexes triggers the ATM/ATR cellular DNA damage checkpoint response.^5-7^ RPA physically interacts with over three dozen DNA processing enzymes and recruits them to the site of DNA metabolism.^4,8-10^ RPA also hands-off the DNA to these enzymes and correctly positions them on appropriate chromosomal structures to facilitate their respective catalytic activity.^11-13^ In addition, several new cell-cycle specific functions have also been recently uncovered. For example, RPA activates a mitosis-specific R-loop driven ATR pathway for faithful segregation of chromosomes.^14^ In concert with RAD52, RPA facilitates mitotic DNA synthesis (MiDAS) to counteract DNA replication stress at common fragile sites loci on the chromosomes.^15,16^

To coordinate such diverse functions, RPA utilizes a unique structural assembly of DNA binding domains (DBDs) and protein interaction domains (PIDs) situated across three subunits - RPA70, RPA32 and RPA14.^17-19^ There are six oligonucleotide/oligosaccharide binding (OB) folds labeled A-F (Figure 1a). Four OB-folds (DBDs - A, B, C and D) contribute most to ssDNA interactions. DBDs-A, B, and C are situated in the large RPA70 subunit and are connected by flexible linkers. DBD-D resides in RPA32. The heterotrimer is held together through extensive physical interactions between DBD-C, DBD-D and the RPA14 subunit (trimerization core; Figure 1a). There are two PIDs; one is OB-F situated at the N-terminus of RPA70 (PID^70N^) and connected to DBD-A through a disordered 80 aa linker. The other PID is a winged helix (wh) domain located at the C-terminus of RPA32 (PID^32C^) and connected to DBD-D by a 34 aa disordered linker. The N-terminal ∼40 aa region in RPA32 is extensively phosphorylated by a slew of kinases (Figure 1a).^20-29^ The flexible linkers allow the domains of RPA to form various configurations (defined as the relative positions of the DBDs and PIDs) and the prevailing hypothesis is that one or more of these configurations drive specific DNA metabolic roles.^4,8,10,11,30-35^ The formation of specific configurations and associated functions are thought to be regulated by post-translational modifications of RPA including phosphorylation, acetylation, sumoylation, and ubiquitination.^4,36^

**Figure 1.**
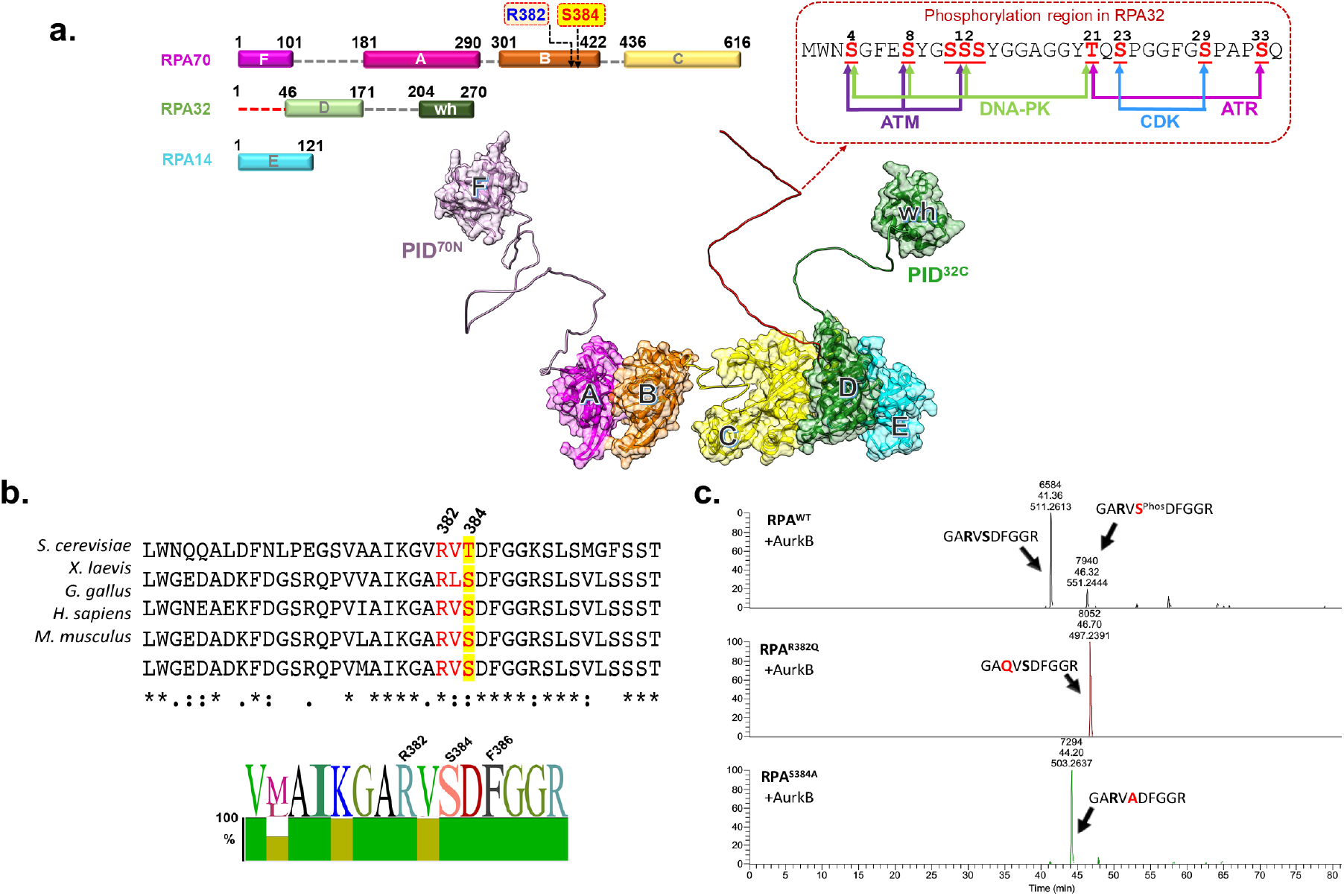
DNA binding domain B (DBD-B) in RPA70 is phosphorylated by Aurora B kinase. **a)** Three RPA subunits RPA70, RPA32, and RPA14 form a heterotrimer and harbor multiple oligosaccharide/oligonucleotide (OB) domains. A, B, C and D are DNA binding domains (DBDs). OB-F and the wh-domain are two protein interaction domains. The N-terminus of RPA32 (shown in red square) is hyperphosphorylated by multiple kinases including ATM, DNA-PK, CDK, and ATR and the known sites of phosphorylation are denoted in the insert. Arg-382 and Ser-384 in the putative Aurora B kinase motif are also noted in RPA70. The structure of human RPA OB-domains and connecting linkers were generated using AlphaFold and aligned to the trimerization core observed in the crystal structure (PDB:1L1O). **b)** Sequence alignment of residues in DBD-B reveal a conserved putative phosphorylation motif for Aurora kinase in higher eukaryotes. A sequence logo representation of the conservation is also shown. **c)** *In vitro* phosphorylation of recombinant RPA with Aurora B, followed by MS-MS analysis, shows residue Ser-384 in RPA70 as the sole site of phosphorylation. Extracted Ion Chromatograms (EICs) for unphosphorylated and phosphorylated tryptic peptide containing S384 in the WT-RPA70 along with EIC of corresponding point mutations are shown. Alanine substitution of the Ser-384 phosphosite, or perturbation of the Aurora B recognition motif through a cancer-associated Gln substitution at Arg-382, results in loss of phosphorylation.

Post-translational modifications of RPA, especially phosphorylation of the RPA32 subunit, has been shown to regulate RPA functions based on the physiological context. RPA is phosphorylated during the S- and mitotic phases by cyclin-dependent protein kinases (CDK1/2)^14,37,38^ and is hyperphosphorylated upon DNA damage by the PI3K-like family of kinases including DNA-PK,^23,39-42^ ATM^20,42-44^ and ATR^7,14,21,42,44-47^. Phosphorylation of RPA32 has been shown to tune the timing and specificity of RPA association with factors involved in DNA replication versus DNA repair.^24,42^ Most studies of RPA phosphorylation have focused on RPA32, and the characterized sites map to a 40-amino acid disordered region in the N-terminus of RPA32 (Figure 1a). Interestingly, phosphorylation in this region modestly influences the DNA binding properties.^28,29,48^ Cells carrying RPA32 lacking the phosphorylation motif are hypersensitive to DNA damaging agents and produce mutator and hyper-mutator phenotypes.^49^ Hyperphosphorylated RPA also inhibits DNA resection by blocking the Blooms helicase.^50,51^ The hyperphosphorylated region of RPA32 is not part of the DBDs or PIDs (Figure 1a). Nevertheless, the precise nature of the domain configurations and how they are modified by hyperphosphorylation remains a mystery.

While phosphorylation of RPA32 is well characterized, and often used as a marker to define replication and repair events, the effects of such modifications on other RPA subunits are not well characterized. In *S. cerevisiae*, Ser-178 in RPA70 is phosphorylated by Mec1^43,52,53^ and we recently showed that a phosphomimetic substitution in yeast RPA promotes cooperative binding of RPA molecules on ssDNA.^36^ In addition, the role of RPA in DNA metabolism in the S-phase has been well characterized, but its function and regulation in mitosis, specifically during an unperturbed phase, are not well defined. Here, we have uncovered a novel mode of regulation of RPA70 that is specific to the mitotic phase of cell cycle. We show that Aurora B kinase phosphorylates Ser-384 in DBD-B of RPA70 in mitosis. The adjacent Arg-382 residue is an integral part of the Aurora B recognition motif and also lies within the binding interface for DSS1-BRCA2.^54^ Biochemically, a RPA^S384D^ phosphomimetic reveals configurational changes in the DBDs and releases the PIDs to promote protein-protein interactions. In addition, interactions with DSS1 are perturbed upon phosphorylation at Ser-384, thus resulting in likely loss of DSS1-BRCA2 recruitment and suppression of homologous recombination during mitosis. Loss of phosphorylation results in increased cell death, activation of p53 checkpoint response, defects in chromosome segregation, and increased sensitivity to DNA damage. Detailed functional and mechanistic characterization reveal an RPA-Aurora B signaling axis that functions through phospho-modification of RPA70 during mitosis and is essential to maintain chromosome segregation fidelity.

## Results

Studies on post-translational modifications of RPA, especially phosphorylation, have focused on the N-terminal region of RPA32 (Figure 1a). Since three of the key DNA binding domains (DBDs-A, B and C) and a major protein interaction domain (PID^70N^) reside in RPA70, knowledge of how post-translational modifications influence these domains and RPA function is essential. Several global phosphoproteomic studies have identified Ser-384 in DBD-B of RPA70 as a prominent site of phosphorylation.^55,56^ Sequence analysis of this region reveals presence of a highly conserved minimal consensus motif [(R/K)_1-3_-X-(S/T)] for Aurora B kinase^57,58^ in eukaryotes (Figure 1b). Furthermore, Arg-382 in this motif has been identified as part of the RPA interaction site for BRCA2-DSS1^54^ and thus warrants detailed functional characterization.

### Aurora B kinase phosphorylates RPA in vitro

To test whether RPA is phosphorylated by Aurora B, we performed *in vitro* kinase assays and analyzed modifications in the RPA subunits using mass spectrometry and autoradiography. Site-specific phosphorylation of RPA70 is observed only in the presence of Aurora B and a single site of phosphorylation at Ser-384 is identified in MS/MS analysis (Figure 1c). Mutation of Ser-384 to Ala results in loss of phosphorylation (Figure 1c and Supplementary Figure 1). Arg-382 in the Aurora B consensus motif is also important as Arg-382 to Gln mutations have been observed in primary skin and thyroid cancers albeit at low frequency (COSMIC).^59^ An Arg-382 to Gln substitution in the motif (RPA^R382Q^) also abolishes phosphorylation by Aurora B as shown by MS analysis and autoradiography (Figure 1c and Supplementary Figure 1). Faint background signal is observed for RPA32 and both mutants of RPA70. However, MS analysis did not identify other sites of phosphorylation in the other two subunits (RPA32 or RPA14) and for RPA70 in the two mutants, under all conditions tested. These data suggest that Aurora B kinase only phosphorylates the RPA70 subunit of RPA at the Ser-384 site.

### RPA phosphorylation by Aurora B is specific to mitosis

Next, we investigated the cellular regulation of RPA by Aurora B. Since Aurora B functions in mitosis, we probed for RPA70 phosphorylation at Ser-384 (pS384-RPA70) in asynchronous and mitotic HCT116 and 293T cells using a custom-generated phospho-specific antibody. In both these cell lines, phosphorylation at the Ser-384 site is observed only during mitosis (Figures 2a & b), which is lost upon entry into G1 phase. A slower migrating form of total RPA70 was also enriched in mitotic cells suggesting phospho-modification (Figure 2a). Induction of Ser-10 phosphorylation of Histone H3 was used as a marker for mitosis. These data show that Aurora B-mediated phosphorylation of RPA is specific to mitosis, in agreement with the well-established mitotic functions of Aurora B.^60^

**Figure 2.**
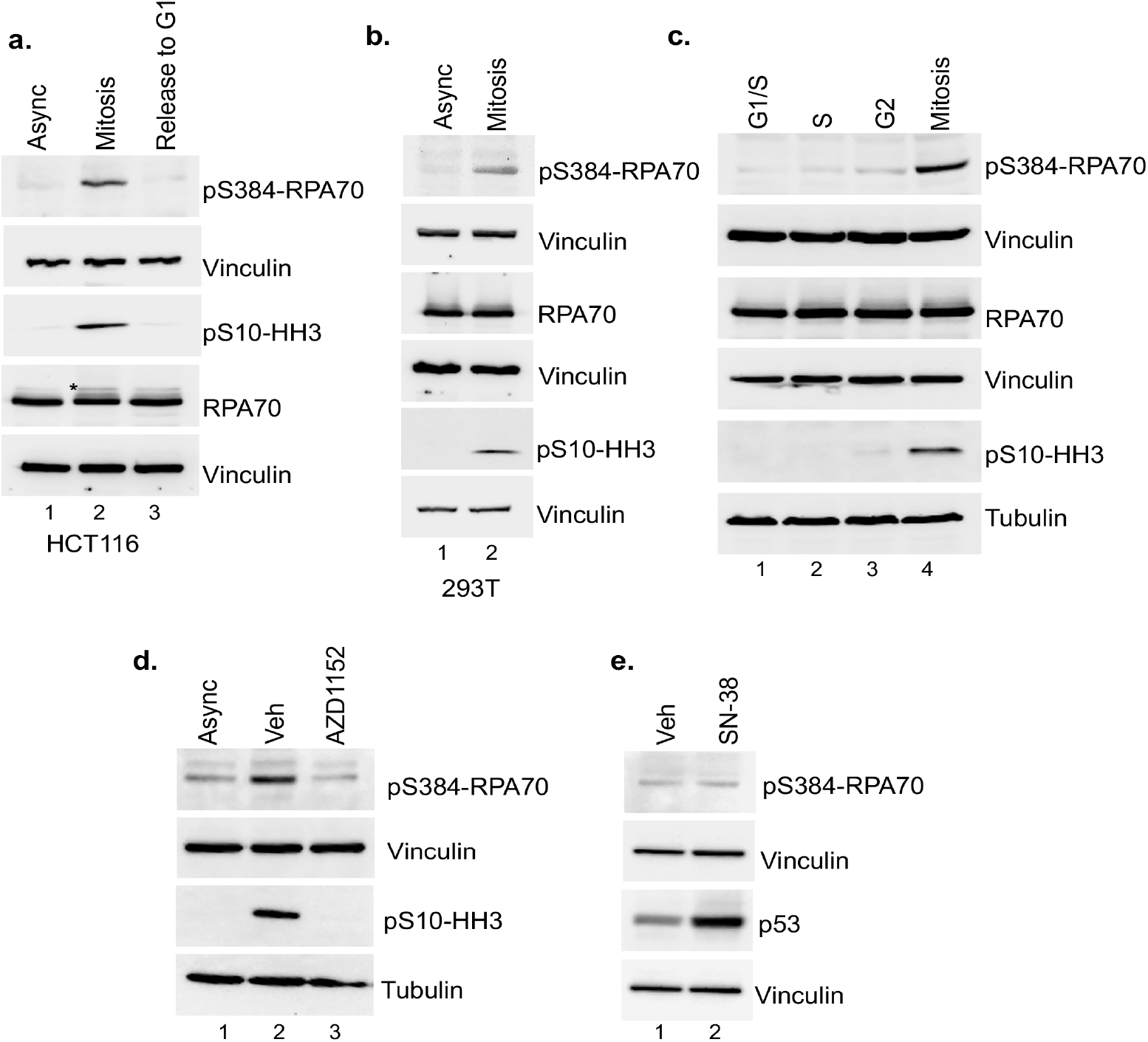
Aurora B phosphorylates RPA at Ser-384 specifically in mitosis. **a)** Western blot analysis of asynchronous (async) HCT116 cells, nocodazole-arrested mitotic cells, and arrested cells released (3hours) into G1 phase. Blots are representative of at least 3 independent replicates. Blots were probed with monoclonal phospho-S384-RPA70 specific (pS384-RPA70) (custom antibody, Genscript) and total RPA70 antibodies. Vinculin was used as a loading control. Mitotic arrest was confirmed using phospho-Ser10-histone H3 specific antibody (pS10-HH3). * Indicates a slower migrating form of RPA70 enriched in mitotic cells. **b)** Similar mitosis specific phosphorylation of RPA70 at Ser-384 was also observed in 293T cells using western blot analysis. All subsequent studies were carried out in HCT116 cells. **c)** RPA70 phosphorylation at Ser-384 was assessed in different phases of the cell cycle by synchronization at G1/S boundary using double thymidine block followed by release into S and G2 phases. Phosphorylation in mitosis was assessed as described in a). Tubulin and vinculin were used as loading controls. Specific phosphorylation of RPA70 at Ser-384 is observed only during mitosis. **d)** Aurora B was selectively inhibited in mitotic cells with short-term treatment (45 min) of 3 mM AZD1152 and results in loss of RPA70 Ser-384 phosphorylation. Inhibition of Aurora B was confirmed by loss of Ser10-Histone H3 phosphorylation. Asynchronous cells and vehicle-treated (0.1 % DMSO) mitotic cells were used as controls. All blots are representative of at least 3 independent replicates. **e)** Cells were treated with vehicle (Veh; 0.1 % DMSO) or 10 ng/ml SN-38 for 21 hours to induce replication stress. DNA damage does not induce RPA70 phosphorylation at Ser-384.

Further assessment of pS384-RPA70 across all phases of the cell cycle by synchronization using double thymidine block show that Aurora B phosphorylation of RPA is observed only in mitosis (Figure 2c). HCT116 cells were efficiently synchronized at the G1/S boundary using double thymidine block as shown by flow cytometry (Supplementary Figure 2). To further determine the specificity of mitotic-specific phosphorylation of RPA70 by Aurora B kinase, cells were treated with an Aurora B-specific inhibitor (AZD1152).^61,62^ Inhibition of Aurora B was confirmed by loss of Ser-10 Histone H3 phosphorylation, which is a well characterized substrate specific for Aurora B in mitosis (Figure 2d). Short-term treatment with the Aurora B inhibitor caused a marked inhibition of Ser-384 phosphorylation of RPA70 suggesting that Aurora B is the kinase that predominantly phosphorylates RPA70 in mitosis at Ser-384. In addition, we found that phosphorylation at Ser-384 is not observed upon induction of replication stress by metabolite of irinotecan (SN-38; Figure 2e). Effect of SN-38 was confirmed by stabilization of p53. These data further showed that phosphorylation at Ser-384 of RPA70 is not regulated by genomic stress.

### Phosphorylation at Ser-384 moderately alters the ssDNA binding properties of RPA

The Aurora B motif in DBD-B of RPA70 is situated within the DNA binding pocket (Figure 3a). In the structure (PDB: 1JMC)^63^, Arg-382 makes several contacts with the backbone of ssDNA. While Ser-384 does not directly interact with the DNA in the structure, it makes backbone interactions with Phe-386, which in turn base-stacks with ssDNA and is thus critical for DBD-B interaction with DNA (Figure 3a). To test if phosphorylation at Ser-384 in RPA70 influences the ssDNA binding properties of RPA, we measured the DNA binding activity of RPA using fluorescein-labeled (dT)_20_ or (dT)_40_ oligonucleotide substrates (Figures 3b & c). In fluorescence anisotropy experiments we see no observable differences in DNA binding activities between RPA and the phosphomimetic mutant RPA^S384D^ (Figures 3b & c). Since RPA has several high-affinity DNA binding domains (DBDs), subtle differences in ssDNA binding by individual DBDs are often hidden by the overall macroscopic DNA binding effect; *i*.*e*., ssDNA binding will be observed irrespective of whether one or more DBDs are bound to DNA.^11^ To better tease out subtle differences in DNA binding properties, we utilized rapid kinetic experiments to capture the rate of RPA binding to ssDNA. Using stopped flow analysis, we monitored the change in intrinsic tryptophan (Trp) fluorescence of RPA upon ssDNA binding (Figures 3d & e). We observed changes in the rate and amplitude of Trp quenching. Plotting the observed rates of Trp fluorescence change versus DNA concentration yields k_on_ and k_off_ for DNA interactions (Figure 3f). RPA^S384D^ has a slower k_on_ (1.9±0.1×10^10^ M^−1^s^−1^) and faster k_off_ (65±7 s^−1^) compared to RPA (k_on_= 5.2±0.3 x10^10^ M^−1^s^−1^ & k_off_ = 39±4 s^−1^) and shows RPA binding to ssDNA with ∼5-fold higher affinity (K_D_= 0.75±0.4 nM) compared to RPA^S384D^ (K_D_ = 3.4±0.2 nM). Introduction of the negative charge at position 384 likely influences DNA binding to DBD-B and/or the path of ssDNA binding within this domain.

**Figure 3.**
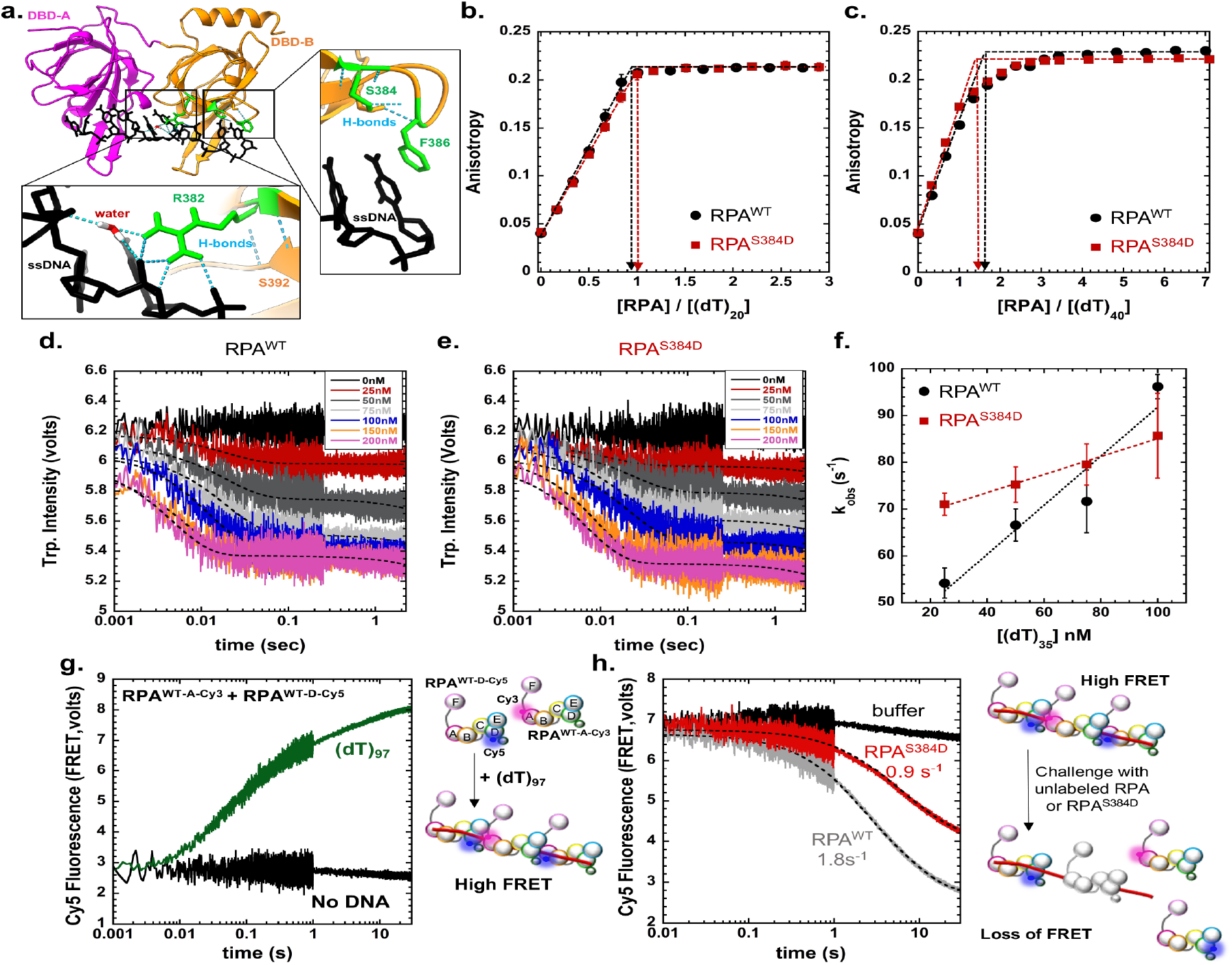
A phosphomimetic S384D substitution minimally influences the DNA binding properties of RPA, but induces configurational changes. **a)** Contacts between R384 and the ssDNA are shown in the crystal structure of DBD-A, B (PDB:1JMC). S384 does not directly contact the ssDNA, but positions F386 to promote a key base stacking interaction with the base. RPA or RPA^S384D^ binding to DNA was measured using anisotropy using either **b)** 5′-FAM-(dT)_20_ or **c)** 5′-FAM-(dT)_40_ ssDNA oligonucleotides. In both cases, high affinity stoichiometric binding is observed for RPA and RPA^S384D^ and no significant changes in the binding behavior is observed. Stopped flow kinetic analysis of ssDNA binding were performed with **d)** RPA or **e)** RPA^S384D^ by monitoring the change in intrinsic tryptophan fluorescence as a function of ssDNA (dT)_35_. **f)** Plot of the k_obs_ versus ssDNA concentration yields k_on_ and k_off_ values. RPA^S384D^ has a slower k_on_ (1.9±0.1×10^10^ M^−1^s^−1^) and faster k_off_ (65±7 s^−1^) compared to RPA^WT^ (k_on_ = 5.2±0.3 x10^10^ M^−1^s^−1^ & k_off_ = 39±4 s^−1^). K_D_ values extracted from these measurements show RPA^WT^ binding to ssDNA with ∼5-fold higher affinity (K_D_ = 0.75±0.4 nM) compared to RPA^S384D^ (K_D_ = 3.4±0.2 nM). **g)** A Förster Resonance Energy Transfer (FRET) experiment was developed using two fluorescent versions of RPA. RPA was site-specifically labeled on either DBD-A with Cy3 or DBD-D with Cy5. Equimolar ratios of both fluorescent RPA were mixed with ssDNA (dT)_97_ in a stopped flow. Changes in Cy5 fluorescence were monitored by exciting Cy3 at 535 nm. Assembly of multiple RPA on the long ssDNA substrate results in a high FRET signal (green trace). In the absence of ssDNA, no enhancement in fluorescence is observed. **h)** RPA filaments were preformed on ssDNA (dT)_97_ using the Cy5- and Cy3-labeled RPA and facilitated exchange activity was measured by challenging the RPA-ssDNA assembly with mixing in unlabeled RPA or RPA^S384D^. RPA exchanges the fluorescent-RPA faster (k_*FE*_ = 1.8 s^−1^) compared to RPA^S384D^ (k_*FE*_ = 0.9 s^−1^).

### Aurora B phosphorylation promotes formation of higher density RPA binding to ssDNA

In the Trp quenching experiments we noticed a significant difference in the basal intrinsic Trp fluorescence between RPA and RPA^S384D^ (Supplementary Figure 3a). Upon ssDNA binding, the amplitude of Trp fluorescence quenching was similar, but the signals did not reach the same plateau. Interestingly, measurement of secondary structure using circular dichroism (CD) shows no difference between RPA and RPA^S384D^ (Supplementary Figure 3b). Thus, the phosphomimetic substitution does not change the folding of DBD-B or other neighboring domains. We hypothesized that the change in intrinsic Trp fluorescence likely originates from altered configuration(s) of the DBDs (orientation/position of DBD-B with respect to the other domains) upon phosphorylation by Aurora B. Changes in RPA configuration promote formation of nucleoprotein filaments that are structurally different. For example, in *S. cerevisiae* RPA, a phosphomimetic substitution at Ser-178 situated just outside of DBD-A promotes cooperative binding of RPA to ssDNA.^36^ Here, the DBD-A from one RPA molecule interacts with DBD-E of the neighboring RPA and these interactions are proposed to be stabilized upon phosphorylation. Since we observe evidence for configurational changes in RPA upon Aurora B phosphorylation at DBD-B, we tested whether binding of multiple RPA molecules on short versus long ssDNA substrates is altered. Binding of RPA or RPA^S384D^ to short (dT)_35_ or longer (dT)_97_ ssDNA oligonucleotides were analyzed using size exclusion chromatography (SEC; Supplementary Figure 4). On the shorter DNA substrate, a single peak corresponding to one RPA bound to ssDNA is observed for RPA (Supplementary Figure 4a). In contrast, for RPA^S384D^, an additional larger species is observed suggesting binding of multiple RPA molecules (Supplementary Figure 4a). This phenomenon is exaggerated on the longer ssDNA substrate where a higher number of RPA^S384D^ molecules are bound compared to RPA (Supplementary Figure 4b). Thus, we propose that Aurora B phosphorylation of RPA is changing the arrangement of DBD-B and this configurational change promotes higher density of RPA molecules bound on ssDNA.

Such configurational changes could affect the stability of the RPA nucleoprotein filament and alter the accessibility of ssDNA. We followed the facilitated exchange (FE) activity of RPA as an experimental measure of filament stability. During FE, RPA bound on ssDNA is replaced by free RPA in solution.^32,64^ While the mechanisms underlying FE are poorly understood, it is thought that the dynamic binding and dissociation of the individual DBDs allow RPA to exchange. Since phosphorylation by Aurora B mildly affects the DNA binding properties of DBD-B, we tested FE on nucleoprotein filaments formed by either RPA or RPA^S384D^. Mixing equimolar concentrations of RPA-DBD-A^Cy3^ and RPA^WT^-DBD-D^Cy5^ with (dT)_97_ ssDNA results in a FRET-induced increase in Cy5 fluorescence when Cy3 is excited (Figures 3g & h). The FRET signal arises from the defined polarity of RPA-ssDNA interactions where DBD-A resides towards the 5′ end of the DNA.^65^ Thus, DBD-A from one RPA molecule sits adjacent to DBD-D of the neighboring RPA. To measure FE, we premixed RPA^WT^-DBD-A^Cy3^, RPA^WT^-DBD-D^Cy5^ and (dT)_97_ and a corresponding high FRET signal is observed (Figure 3g). When this filament is challenged with either unlabeled RPA or RPA^S384D^, loss of the FRET signal is observed as the fluorescent RPA molecules are replaced by the unlabeled RPA or RPA^S384D^ during FE.RPA is more efficient in performing FE (k_*FE*_ = 1.8 s^−1^) compared to the phosphomimic form RPA^S384D^ (k_*FE*_ = 0.9 s^−1^; Figure 3h). Thus, phosphorylation at Ser-384 and the resulting change in configuration of DBD-B enables easier remodeling of RPA nucleoprotein filaments.

### Loss of Ser-384 phosphorylation of RPA70 markedly affects cell viability

To determine the physiological significance of configurational changes in DBD-B induced by Ser-384 phosphorylation, we generated a homozygous knock-in of Ser-384-Ala RPA70 mutant in the endogenous loci in HCT116 cells. As expected, the phospho-dead mutant (*RPA*^*SA/SA*^) does not show phosphorylation at the Ser-384 site in mitosis (Figure 4a). This finding further validates the specificity of the antibody used. In addition, the phospho-dead mutation does not affect the endogenous levels of total RPA70 (Figure 4a). To investigate the functional effects of Ser-384 phosphorylation, we first investigated cell viability using MTS assays. Cell viability is markedly reduced in the *RPA*^*SA/SA*^ mutant relative to the isogenic parental wild type (*RPA*^*WT/WT*^) cells (Figure 4b), which is consistent with morphological assessments that indicate increased cell death in the *RPA*^*SA/SA*^ mutant. We also observed similar loss of viability in a second clone of *RPA*^*SA/SA*^ mutant (Supplementary Figure 5). Significantly higher rates of apoptosis in the *RPA*^*SA/SA*^ mutant were further confirmed by measuring Caspase-3 and Caspase-7 enzymatic activity using the Caspase-3/7 glo assay (Figure 4c). The phospho-dead mutant was also subjected to replication stress induced by SN-38 (Figure 4d) and showed enhanced sensitivity to replication stress (Figure 4e).

**Figure 4.**
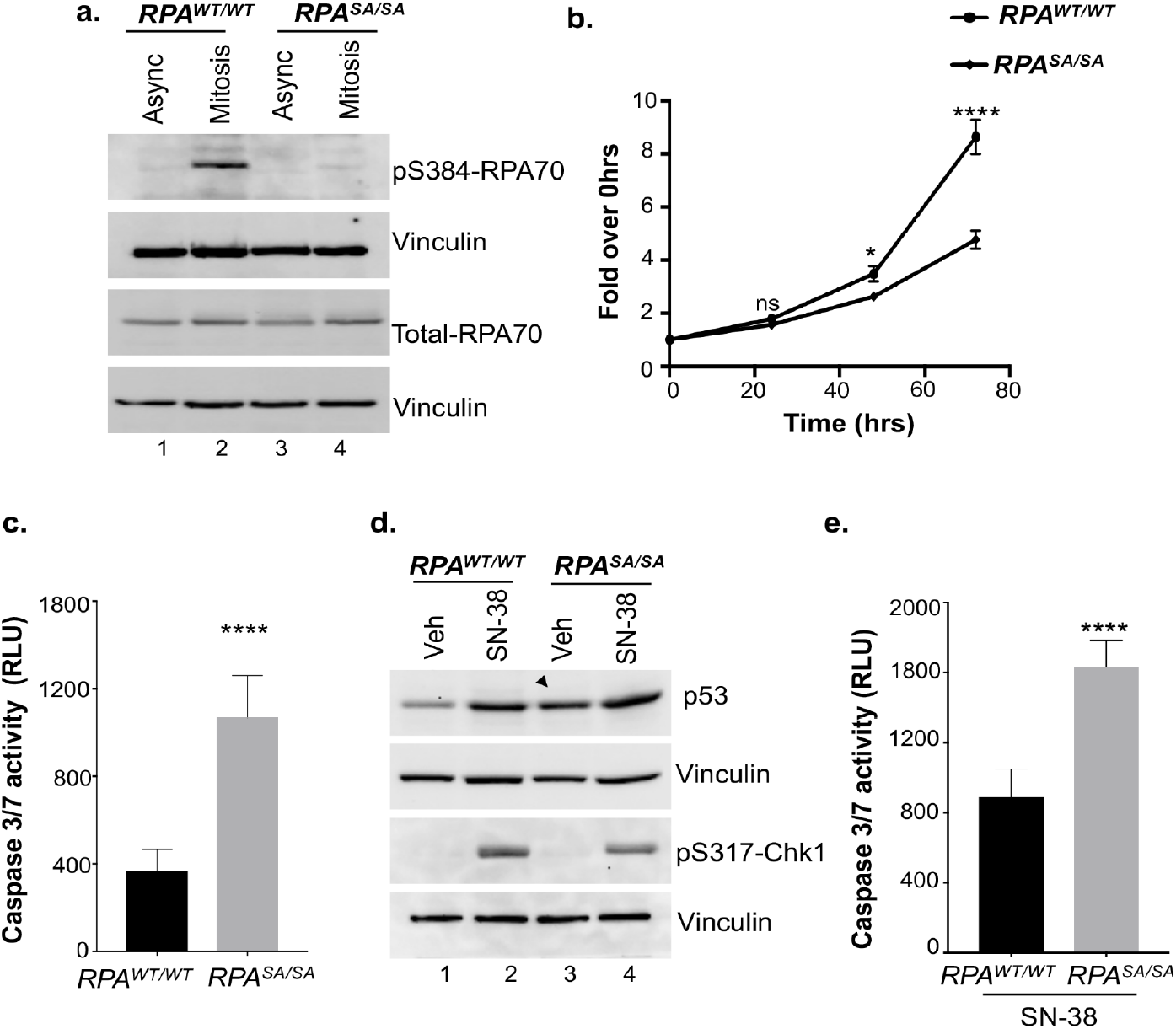
Loss of Ser384-RPA70 phosphorylation markedly disrupts cell viability and induces basal genomic stress response. **a)** Homozygous knock-in of Ser-384-Ala RPA70 phospho-dead mutant in HCT116 cells was generated using CRISPR-Cas9 editing. Cells were synchronized in mitosis using nocodazole and representative western blot depicts loss of RPA70-Ser-384 phosphorylation in RPA^*SA/SA*^ mutant. Asynchronous parental HCT116 cells and cells arrested in mitosis were used as controls. Blots were probed with the indicated antibodies. Blots are representative of at least three independent experiments. **b**) MTS assay shows decreased viability of phospho-dead *RPA*^*SA/SA*^ mutant. Cells were assayed at 0, 24, 48 and 72 hours of growth. Values corrected for background absorbance were normalized to 0hrs of growth. Error=SEM. Mean of three independent experiments were plotted. Triplicate wells were assayed per time point for each experiment. Statistical significance was determined using an unpaired two-tailed *t*-test: \**p*=0.014 at 48 hours, **** *p*<0.0001 at 72 hours, ns=not significant. **c**) Caspase 3/7 activity was significantly higher in the RPA mutant cells as determined by the Caspase3/7 glo assay. Bar graph depicts data corrected for background absorbance. Error=SEM. Mean of three independent experiments were plotted. Triplicate wells were assayed per experiment. Statistical significance was determined using an unpaired two-tailed *t*-test: **** *p*<0.0001. **d**) Representative western blot shows replication stress-response in cells treated with vehicle (veh, 0.1% DMSO) or 20ng/mL SN-38 for 90 min. Arrow indicates increased basal genomic stress as shown by high levels of p53 in the *RPA*^*SA/SA*^ mutant. Data are representative of three independent experiments. **e**) Caspase3/7 activity was determined in response to replication stress by treating cells with 10ng/mL SN-38 for 21 hours and by using Caspase3/7 glo assay. Bar graph displays data corrected for background absorbance. Error=SEM. Mean of three independent experiments were plotted. Triplicate wells were assayed per experiment. Statistical significance was determined using an unpaired two-tailed *t*-test: **** *p*<0.0001.

Analysis of asynchronous cells did not reveal any differences in the percentage of S-phase cells that could account for increased sensitivity of the phospho-dead mutant to replication stress. A 3% to 4% decrease in G1 population of the phospho-dead mutant was observed (Supplementary Figures 6a-c). Interestingly, the phospho-dead mutant displayed high basal levels of p53 even in the absence of SN-38 treatment indicating that the mutant cells are in a constant state of genomic stress that leads to enhanced p53 stability (Supplementary Figure 6d). Mild increase in γH2AX levels were also observed in the mutant cells (Supplementary Figure 6d). However, unlike the p53 checkpoint response, loss of phosphorylation does not activate the Chk1-mediated checkpoint response at a basal state (Figure 4d). In addition, the p53 and Chk1-mediated checkpoint responses were induced by replication stress to a similar extent between the WT and mutant cells (Figure 4d). This indicates that the DNA damage-induced checkpoint response is not altered by loss of Ser-384 phosphorylation. However, loss of phosphorylation enhances basal genomic stress-response and induces extensive apoptosis that further sensitizes cells to replication stress.

### Mitotic phosphorylation of RPA70 is critical for chromosome segregation

Since Ser-384 phosphorylation of RPA70 occurs specifically in mitosis, we wanted to test if defects in mitotic progression contribute to genomic instability. Mitosis is defined by the precise segregation of sister chromatids to opposite spindle poles. Defects in chromosome segregation and nondisjunction can lead to marked increase in anaphase DNA bridges between the segregating sister chromatids.^66,67^ To uncover the functional significance of mitosis-specific phosphorylation of RPA70, we assessed the mitotic phases of the phospho-dead mutant. Interestingly, both in unperturbed mitosis in asynchronous populations and in cells released from pro-metaphase arrest, there was a marked increase in chromosome segregation defects induced by loss of phosphorylation (Figure 5 and Supplementary Figure 7a). Segregation defects were highlighted by the enhanced presence of anaphase bridges and lagging chromosomes. We did not observe enhanced defects in spindle pole formation or spindle alignments in the phospho-dead mutant relative to WT, which indicated that the chromosome segregation defects are likely driven by unresolved replication or recombination intermediates or misalignment of chromatid kinetochore and microtubule attachments. Intriguingly, when we probed for Ser-10 phosphorylation of Histone H3 (the mitotic marker), we observed a 50% decrease in Ser-10 phosphorylation in the phospho-dead mutant (Figures 6a & b). This change in phosphorylation of Histone H3 could not be attributed to changes in total Histone H3 levels or due to differences in mitotic synchronization as shown by flow cytometric analysis (Figures 6c-e). Since phosphorylation of Ser-10 in Histone H3 is important for chromosome condensation, although the mechanism remains poorly resolved, we compared chromosome compaction in WT and phospho-dead mutant as determined by surface area of the nuclei (Supplementary Figure 7b). Consistent with the decrease in Ser10-phosphorylation of Histone H3, there was less chromosome compaction of many nuclei in the phospho-dead mutant (Figure 6f). Thus, these chromosomal defects clearly indicate that Ser-384 phosphorylation of RPA70 is important for segregation of chromosomes in mitosis.

**Figure 5.**
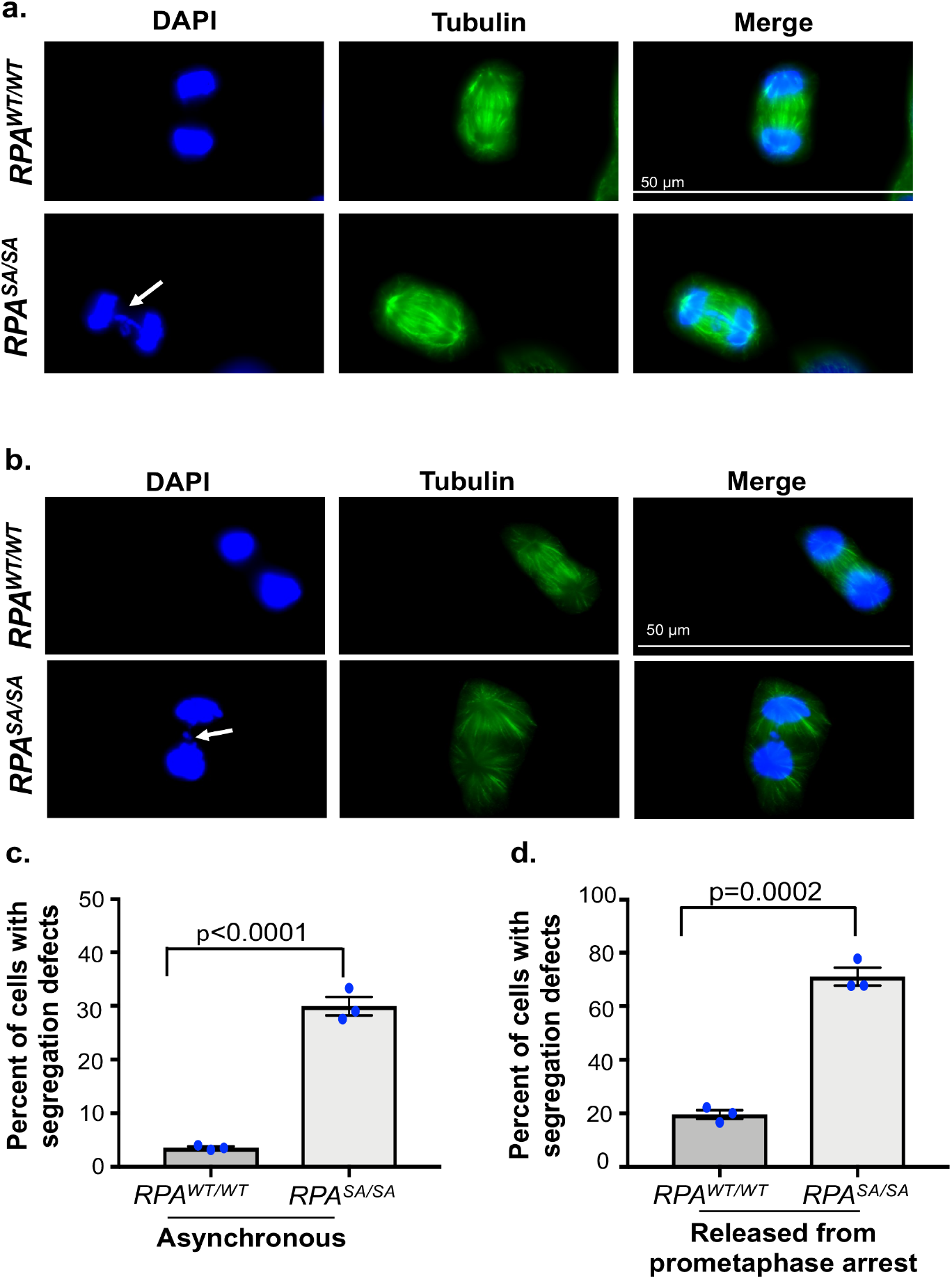
Defects in chromosome segregation induced by loss of Ser384-RPA70 phosphorylation. **a and b)** Representative immunofluorescent images stained with DAPI and anti-Tubulin antibody depict anaphase bridges (white arrow) in the mitotic population of asynchronous *RPA*^*SA/SA*^ mutant and *RPA*^*WT/WT*^ cells (**a**) and in cells released from prometaphase arrest (**b**). Images are representative of three independent experiments. **c** and **d**) Quantitation of anaphase bridges and lagging chromosomes in mitotic population of asynchronous cells (**c**) and in cells released from prometaphase arrest (**d**) *RPA*^*WT/WT*^ and *RPA*^*SA/SA*^ cells. More than 70 cells were counted from three independent experiments. Data represent SEM and statistical significance was determined using an unpaired two-tailed *t*-test.

**Figure 6.**
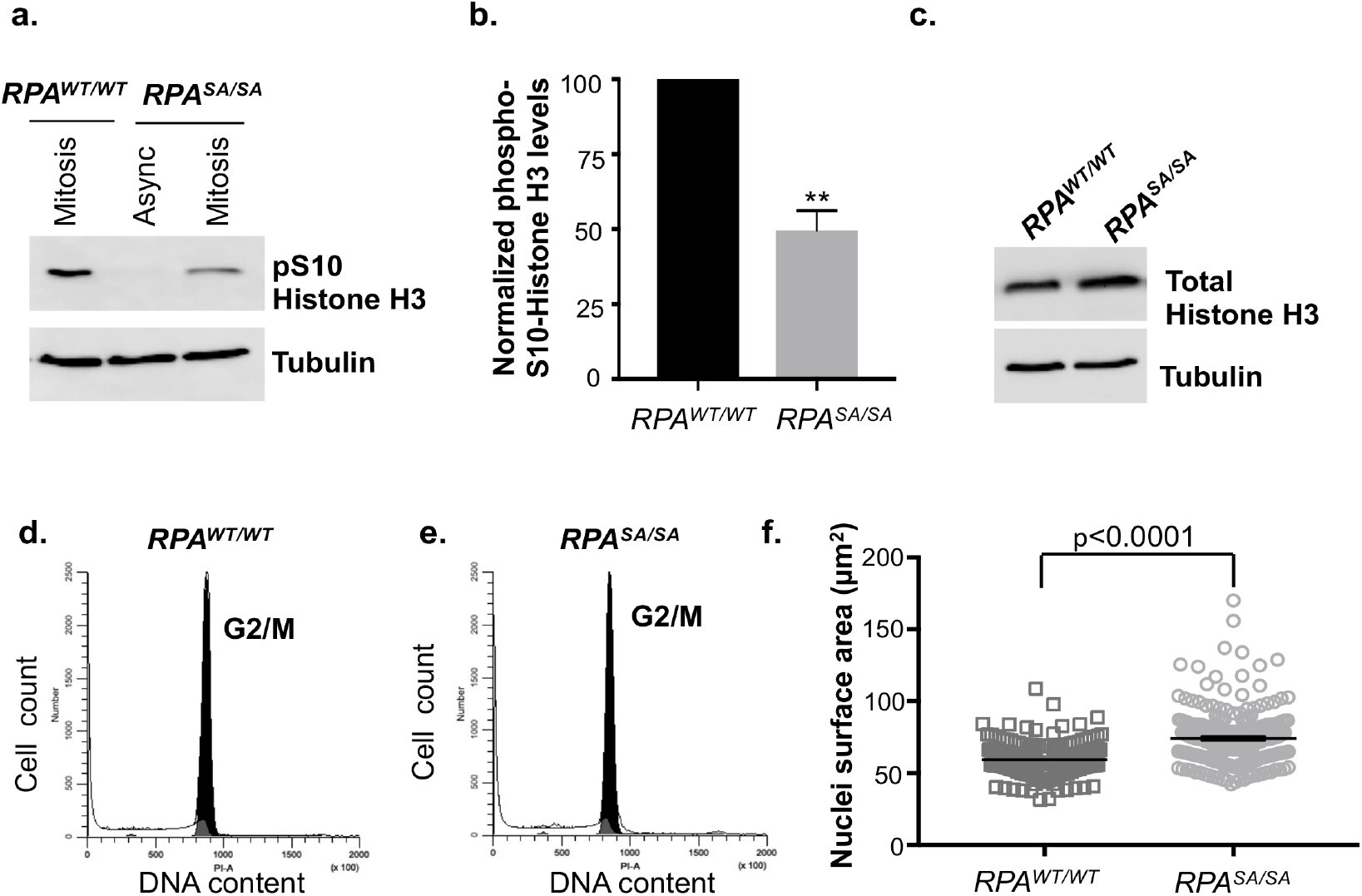
The phoshodead *RPA*^*SA/SA*^ mutant exhibits reduced phosphorylation of Histone H3. **a**) Western blot represents decrease in Ser10 Histone H3 phosphorylation in *RPA*^*SA/SA*^ mutant relative to wild type cells. Asynchronous (asynch) cells were assayed as control. Blots are representative of three independent experiments. **b**) Blots represented in a) were quantitated and normalized to loading control (Tubulin). Data are expressed as a fold over *RPA*^*WT/WT*^. Data are presented as mean of three independent experiments and error=SEM. Statistical significance was determined using an unpaired two-tailed *t*-test: **p=0.0016. **c**) Western blot represents total Histone H3 levels in *RPA*^*SA/SA*^ mutant and parental cells. Blots are representative of three independent experiments. **d and e**) *RPA*^*WT/WT*^ (**d**) cells synchronized in mitosis and collected by shake-off method were found to be arrested in G2/M phase of cell cycle similar to *RPA*^*SA/SA*^ mutant (**e**) as analyzed by flow cytometry. DNA content was determined using propidium iodide staining. Data represents three independent experiments. **f**) Surface area of nuclei stained for phospho Ser-10 Histone H3 represented in supplementary figure 7b. were quantitated using Image J analysis. Data is presented as a scatter plot of more than 200 nuclei measured across three independent measurements. Error=SEM. Statistical significance was determined using an unpaired two-tailed *t*-test.

### Phosphorylation by Aurora B changes the configuration of RPA domains

To better understand how phosphorylation by Aurora B influences the configurational changes within the multi-domain structure of RPA, we performed hydrogen-deuterium exchange mass spectrometry (HDX-MS) analysis of RPA^WT^ and RPA^S384D^ in the absence or presence of ssDNA (Figure 7). Configurational changes bring about changes in uptake or loss of deuterium^30^, and these HDX changes (uptake or loss) are plotted as a comparison between RPA and RPA^S384D^ (Figure 7, and Supplementary Figures 9 -15). These changes were further assessed in the absence or presence of ssDNA. Surprisingly, changes in HDX are observed in several regions including DBDs-A, B, the trimerization core, the two protein interaction domains, and the flexible linker between DBD-B and DBD-C (Figure 7a). In the presence of ssDNA, the patterns of HDX change (Figure 7b). However, the changes again cover multiple domains including DBDs-A, B, the trimerization core, and one of the protein interaction domains (OB-F; Figure 7b). These changes persist over longer time scales suggesting that the configurational changes driven by the phosphomimetic substitution are stable (Supplementary Figure 8). These data reveal that the various domains of human RPA do not exist in a splayed-out fashion, but likely interact with each other in a configurationally compacted form. These interactions are altered upon phosphorylation at Ser-384. Since the DNA binding properties are not severely affected, another functional outcome could be the modulation of protein-protein interactions between RPA and RIPs. Since OB-F in RPA70 (PID^70N^) and the winged-helix domain in RPA32 (PID^32C^) show changes in HDX (Figures 7a & b), phosphorylation at Ser-384 also likely releases them and promotes RPA-protein interactions through these domains.

**Figure 7.**
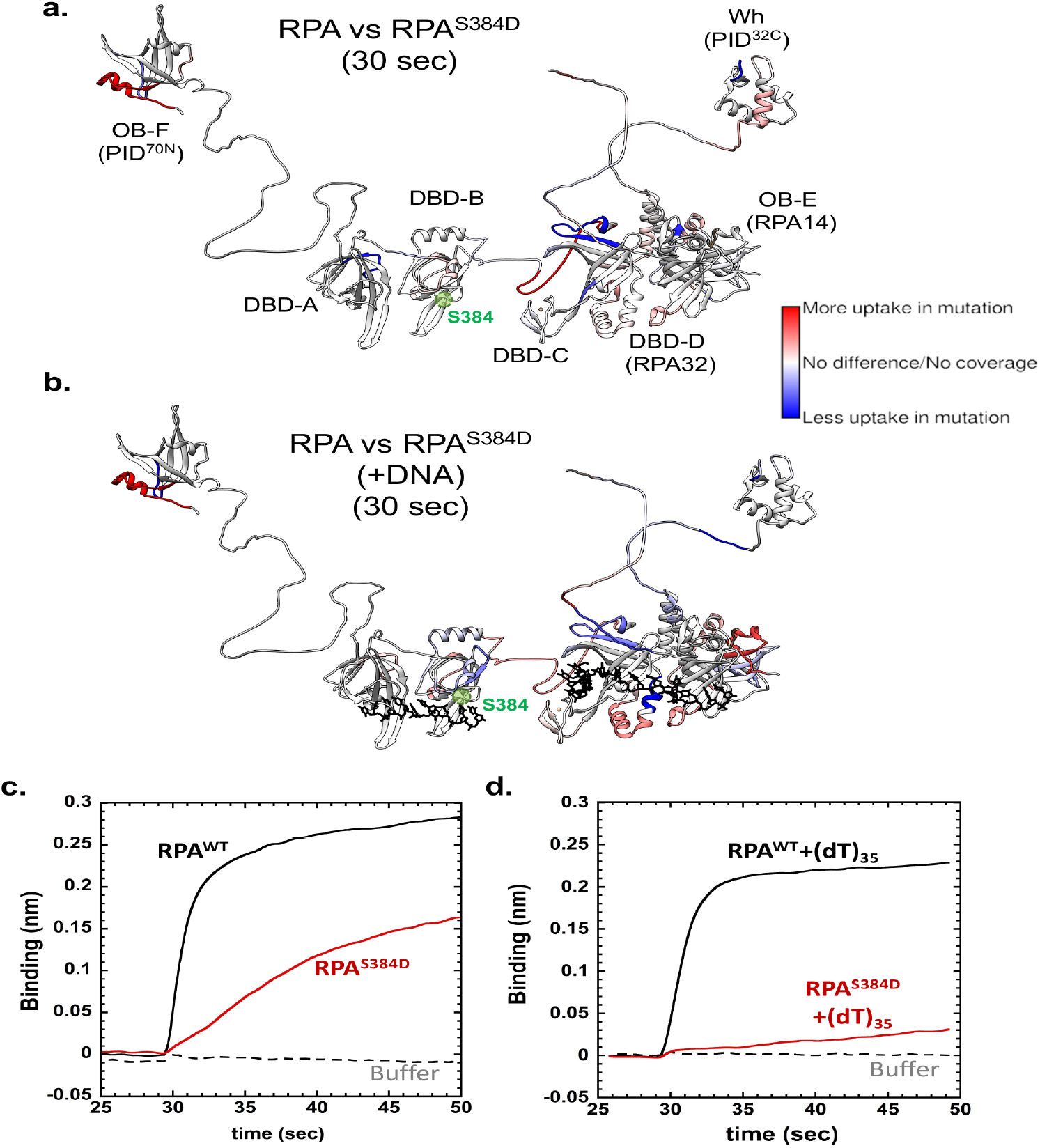
Configurational changes in RPA are induced by a S384D substitution within the Aurora B motif in DBD-B. HDX changes between RPA and RPA^S384D^ are shown in the **a)** absence or **b)** presence of ssDNA (ssDNA is depicted in black). Changes in deuterium uptake/loss are observed in almost all DNA binding and protein-interaction domains. Data are mapped onto the structure of human RPA which is built using the structures of the OB domains from crystal structures. The flexible linkers were modeled using AlphaFold. Position of Ser-384 is denoted in green. Bio-layer interferometry analysis of RPA or RPA^S384D^ binding to DSS1 in the **c)** absence or **d)** presence of ssDNA. RPA^S384D^ shows reduced binding to DSS1 in the absence of DNA. When ssDNA-bound, almost complete loss of DSS1 binding to RPA is observed.

### Ser-384 phosphorylation inhibits DSS1 binding to RPA

While PID^70N^ and PID^32C^ primarily coordinate most of the characterized RPA interactions with RIPs, a few interactors bind to other regions in RPA70. For example, physical interactions between DSS1 and RPA have been mapped to DBD-B.^54^ DSS1 works in complex with BRCA2 to facilitate the loading of RAD51 on RPA-coated ssDNA.^68^ Physical interaction between DSS1 and RPA is required for this activity and the binding interface resides within the F-A-B half of RPA. Interestingly, in NMR studies, Arg-382 is one of the residues that show a chemical shift perturbation upon DSS1 binding.^54^ Arg-382 is part of the Aurora B kinase motif (Figure 1b) and is found mutated in certain cancers. Thus, we tested whether phosphorylation in this motif interferes with DSS1 binding using biolayer interferometry. Strep-tagged DSS1 was tethered onto a streptavidin-coupled optical probe and binding to RPA^WT^ or RPA^S384D^ was measured. A reduction in binding to DSS1 is observed for RPA^S384D^ (Figure 7c). Since DSS1-BRCA2 would encounter RPA-ssDNA structures in the cell, we tested the binding interaction when RPA is in complex with ssDNA. In this context, complete loss of binding to DSS1 is observed for RPA^S384D^ (Figure 7d). Thus, Ser-384 phosphorylation by Aurora B perturbs RPA binding to DSS1 likely resulting in the inhibition of BRCA2 recruitment and suppression of HR in mitosis.

## Discussion

Maintenance of genomic integrity depends on the faithful segregation of newly replicated genomic DNA to the daughter cells during mitosis. RPA coordinates almost all aspects of DNA metabolism and functions by binding to ssDNA and recruiting over three dozen enzymes. In the S-phase, RPA coordinates DNA replication, repair, and recombination. However, in mitosis, protein-protein interactions of RPA and its role in chromosomal repair and recombination has remained poorly understood. How RPA gains functional specificity for specific events in various parts of the cell cycle and tailor’s ssDNA handoff to other enzymes is also a long-standing question. Modulation of RPA activity through post-translational modifications have been characterized in the S-phase checkpoints where kinases such as ATM/ATR are recruited onto RPA-coated ssDNA and phosphorylate RPA32 to activate appropriate checkpoints.^6,42^ Furthermore, RPA32 phosphorylation in mitosis by CDK1 promotes mitotic exit in the presence of DNA damage.^20^ However, disruption of CDK1-specific mitotic phosphorylation of RPA does not affect normal mitotic progression.^69^ Therefore, how RPA activity in mitosis is regulated under normal unstressed conditions and in the absence of DNA damage is not known.

Here, we uncover site-specific phosphorylation of RPA by the mitosis-specific Aurora B kinase in the large subunit (RPA70) at Ser-384 within DNA binding domain B (DBD-B). Aurora B kinase is a key component of the chromosomal passenger complex. It contributes to numerous processes that maintain the fidelity of cell division including kinetochore stabilization, kinetochore-microtubule attachment, spindle assembly checkpoint and abscission checkpoint.^70^ Aurora B coordinates a complex network of cellular interactions that are tied to its state of activation and localization on the centromeres.^71,72^ Interestingly, RPA also localizes to the centromeres and binds to ssDNA associated with R-loops.^14^ Thus, the spatial regulation of RPA by Aurora B at the centromeres could enable sensing of ssDNA during mitosis and protect ssDNA generated by centromeric stress. Therefore, it is likely that the disruption of this signaling axis disrupts proper chromosome segregation and leads to extensive apoptosis through basal activation of the p53-mediated checkpoint response. Previous studies have shown that delays in mitotic progression can induce p53-dependent cell death response.^73,74^

In the crystal structures of RPA bound to DNA, Ser-384 (the site of Aurora B phosphorylation) does not directly contact ssDNA. Through hydrogen-bonding interactions it positions a neighboring conserved Phe-386 that base-stacks with ssDNA (Figure 3a). Furthermore, Arg-382 which is a part of the Aurora B recognition motif, interacts with the ssDNA backbone. Thus, upon phosphorylation, this pocket in DBD-B is likely remodeled. While phosphorylation does not affect overall DNA binding activity, higher density of RPA^S384D^ molecules bind to ssDNA. The structural effects are more striking in the overall configurational changes of the domains. PID^70N^ (OB-F) and PID^32C^ (winged-helix domain) are two protein-protein interaction domains situated in RPA70 and RPA32, respectively (Figure 1a). Conventionally, these PIDs are assumed to be exposed and readily accessible for binding to RPA interacting proteins (RIPs).

In yeast RPA, we recently showed that Rtt105 (a chaperone-like protein) binds and configurationally compacts the many domains of RPA, including PID^70N^ and PID^32C^.^75^ This phenomenon, termed ‘configurational stapling’, enables Rtt105 to block binding of RIPs (such as Rad52) to RPA. Presence of ssDNA licenses RPA-RIP interactions by remodeling Rtt105. Based on the comparative HDX-MS analysis, we here propose that human RPA has evolved to resemble configurational stapling without the need for a chaperone-regulated mechanism under certain physiological contexts. HDX changes are observed in both PID^70N^ and PID^32C^ when data from RPA and RPA^S384D^ are compared (Figure 7). These data suggest that these PIDs that are connected to their respective DBDs through flexible linkers are not extended out, but likely in close proximity to the other DBDs. This occlusion likely prevents them from binding to RIPs. Upon phosphorylation by Aurora B, the PIDs are ‘licensed’ to bind to select RIPs; in this case, promotion of protein-interactions with mitosis-specific proteins (Figure 8). For example, RPA enriched from mitotic cells were shown to exhibit reduced physical interactions with ATM, DNA pol-alpha, and DNA-PK and treatment with phosphatase restored these interactions.^29^ The enhancement in intrinsic Trp fluorescence in the RPA^S384D^ variant further supports a model for release of configurational compaction upon phosphorylation. The compaction and release of RPA could serve as a potential mechanism to regulate RPA-protein interactions in the cell. Such configurations can be further modulated by additional post-translational modifications.^76^

RPA-RIP interactions can be positively or negatively regulated by phosphorylation.^6,29^ An example of negative regulation is the inhibition of DSS1-BRCA2 binding to the phosphomimetic RPA^S384D^. DSS1-BRCA2 recruitment to ssDNA by RPA is important for resolution of ssDNA intermediates through HR.^54^ Intriguingly, HR is suppressed during mitosis.^77^ While there are several mechanisms that could contribute to HR suppression in mitosis including the inaccessibility of compacted sister chromatids, we propose that inhibition of BRCA2 recruitment during mitosis provides an additional mechanism for HR suppression in mitosis. DSS1, is a BRCA2 partner, and facilitates the RPA-BRCA2 interaction. The mapped region of interaction between DSS1 and RPA resides around Arg-382 in DBD-B and our findings show that Aurora B phosphorylation at Ser-384 blocks DSS1 binding to RPA that will suppress HR in mitosis. The importance of suppressing BRCA2 activity in mitosis is further highlighted by the phosphorylation of BRCA2 by CDKs at Ser-3291. This modification in BRCA2 blocks physical interaction with RAD51. In the S and G2 phases, phosphorylation at Ser-3291 is low, but drastically increases in mitosis.^77^ Thus, RAD51 promoted HR is likely suppressed in mitosis upon phosphorylation of BRCA2. Thus, in addition to CDK regulation of BRCA2, Aurora B phosphorylation of RPA provides added stringency to HR suppression in mitosis. The significance of the DSS1 interaction region is further highlighted by the mutation of the Arg-382 site in certain cancers suggesting that deregulation of this region could underlie genomic instability.

**Figure 8.**
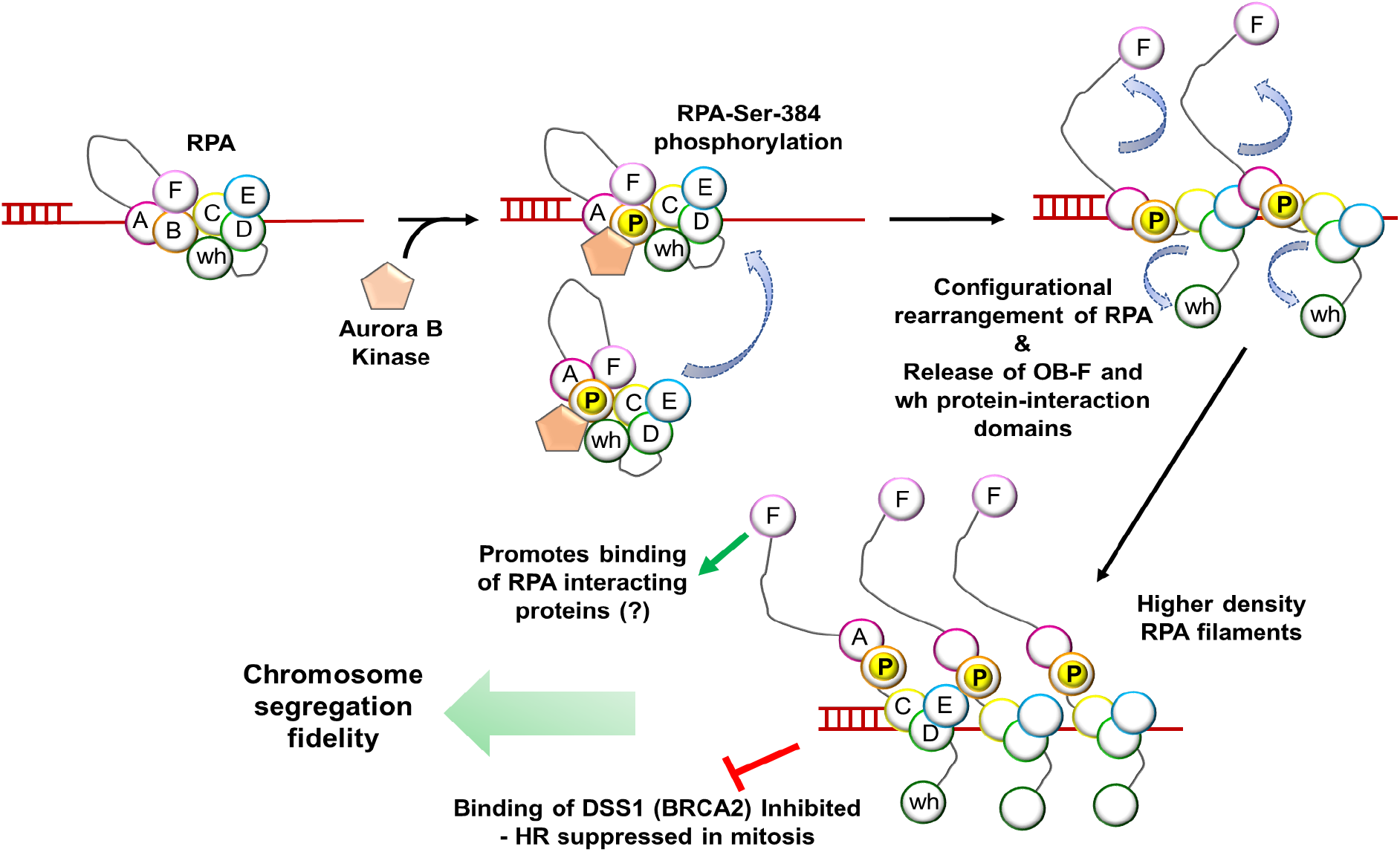
Model for the Aurora B-RPA signaling axis. During mitosis, RPA-bound to ssDNA intermediates or free in solution are phosphorylated by Aurora B kinase at Ser-384 (DBD-B) in the large RPA70 subunit. We propose that phosphorylation releases the protein-interaction domains (OB-F or PID^70N^ and wh or PID^32C^) and promotes formation of higher-density RPA-bound ssDNA filaments. The protein-interaction domains can then promote binding to RPA-interacting proteins. Since the site of phosphorylation resides close to the DSS1 binding site, recruitment of DSS1-BRCA2 is inhibited leading to suppression of homologous recombination during mitosis. Deregulation of the Aurora B-RPA signaling axis leads to errors in chromosome segregation fidelity.

Another interesting observation is the significant reduction of histone H3 phosphorylation and modest compaction of chromosomes in the RPA^*SA/SA*^ cells. The mechanistic basis of these observations, driven by a lack of RPA phosphorylation by Aurora B, is not clear. However, RPA has been shown to directly interact with histone subunits H3-H4^78^ and various histone chaperones including HIRA^79^, CAF-1^78^, and FACT.^80^ Functionally, these interactions promote histone deposition to replication intermediates, H3.3 binding at promoter and enhancer regions, and the recruitment of transcription factors such as Heat Shock Factor-1 to nucleosome-coated DNA. Recent studies have also shown that recruitment of the ubiquitin E3 ligase Bre1 to promote monoubiquitination of histone H2B is driven by physical interactions between RPA and Bre1.^81^ Whether such RPA-regulated histone properties are related to roles in mitosis remains to be investigated.

Under conditions where DNA damage is encountered in mitosis, RPA serves as a cellular marker to define anaphase bridges, ultrafine bridges, and as loci-markers for DNA repair events. Recruitment of the Plk1-interacting checkpoint helicase (PICH) chromatin remodeler and the Bloom’s syndrome helicase (BLM) helicase limit assembly of histones with centromeric DNA during anaphase.^50,82,83^ The activity of both these helicases is required to promote proper resolution of the chromosomes in mitosis. BLM directly interacts with RPA through interactions with PID^70^ the binding is required to stimulate unwinding activity of BLM.^84^ PICH binds to BLM through its C-terminal region. Interestingly, acidic motifs are found in RPA interacting proteins that interact with PID^70N^ of RPA, and several such potential motifs are found in PICH. We show here that Aurora B phosphorylation releases PID^70N^ and likely facilitates interactions with RIPs that bind to this domain (Figure 8). It will be interesting to test whether PICH directly interacts with RPA to function as a complex with BLM during mitosis.

In summary, we propose that Aurora B phosphorylation of RPA acts as a signaling axis that is important for faithful chromosome segregation in mitosis. Thus, future studies of RPA in DNA metabolism should also consider post-translational modifications (especially phosphorylation) at subunits other than RPA32. Tandem modifications at both RPA70 and RPA32 could further yield an additional layer of regulation that might be important in understanding how RPA imparts specificity to DNA repair processes and the maintenance of genomic stability.

## Methods

### Recombinant overproduction and purification of RPA

Human RPA (RPA) was recombinantly expressed using plasmid p11d-hRPA-WT (a kind gift from Marc Wold, Univ. of Iowa). RPA^R382Q^, RPA^S384D^, and RPA^S384A^ mutants were generated in this plasmid background using Q5 site-directed mutagenesis (New England Biolabs). RPA, RPA^R382Q^, RPA^S384D^, and RPA^S384A^ were purified from *E. coli* as described^85^ with minor modifications. Briefly, the appropriate plasmids were transformed into Rosetta2(DE3)pLysS cells (Novagen) and transformants on Luria Broth (LB) were selected using ampicillin (100 μg/ml). An overnight culture (10 ml) from a single colony was grown and added to 1 L of LB media containing ampicillin. Cells were grown at 37 °C until the OD_600_ reached 0.6 and then induced with 0.3 mM isopropyl-b-D-1-thiogalactoside (IPTG). Induction was carried out at 37 °C for 3 hours and the harvested cells were resuspended in 120 ml cell resuspension buffer (30 mM HEPES, pH 7.8, 300 mM KCl, 0.02% Tween-20, 1.5X protease inhibitor cocktail, 1 mM PMSF, 10% (v/v) glycerol and 10 mM imidazole). Cells were lysed using 0.4 mg/ml lysozyme followed by sonication. Clarified lysates were fractionated on a Ni^2+^-NTA agarose column (Gold Biotechnology). RPA was eluted using cell resuspension buffer containing 400 mM imidazole. Fractions containing RPA were pooled and diluted with H_0_ buffer (30 mM HEPES, pH 7.8, 0.02% Tween-20, 1.5X protease inhibitor cocktail, 10% (v/v) glycerol and 0.25 mM EDTA pH 8.0) to match the conductivity of buffer H_100_ (H_0_ + 100 mM KCl), and further fractionated over a fast-flow Heparin column (Cytiva). RPA was eluted using a linear KCl gradient H_100_–H_1500_, and fractions containing RPA were pooled and concentrated using an Amicon spin concentrator (30 kDa molecular weight cut-off; Millipore Sigma). The concentrated RPA was next loaded onto a HiLoad Superdex S200 column (Cytiva) and fractionated using RPA storage buffer (30 mM HEPES, pH 7.8, 300 mM KCl, 0.25 mM EDTA, 0.02% Tween-20, and 10% (v/v) glycerol). Purified RPA protein was flash frozen using liquid nitrogen and stored at −70 °C. RPA concentration was measured spectroscopically using ε_280_ = 87,210 M^−1^cm^−1^.

### Generation of fluorescent hRPA

Fluorescently labeled human RPA with Cy3/Cy5 positioned on DBD-A (RPA-DBD-A^f^) or DBD-D (RPA-DBD-D^f^) were generated using non-canonical amino acids (ncAA) as described for *S. cerevisiae* RPA.^86^ Briefly, 4-azido-*L*-phenylalanine (4AZP) was site-specifically incorporated into DBD-A (RPA70 Ser-215) or DBD-D (RPA32 Trp-107) by engineering TAG stop codons at the corresponding positions in the plasmid using site directed mutagenesis. A 6x-poly-histidine affinity tag was also engineered at the C-terminus of RPA70 or RPA32 to isolate RPA-DBD-A^4AZP^ or RPA-DBD-D^4AZP^ from the truncated non-4AZP incorporated proteins, respectively. The respective plasmid was cotransformed into *E. coli* BL21 Rosetta2(DE3)pLysS cells with a pDuel2-pCNF plasmid that codes for the orthogonal tRNA^UAG^ and tRNA synthetase for 4AZP incorporation.^32^ Cotransformants were selected using both ampicillin (100 mg/ml) and spectinomycin (50 mg/ml). A 10 ml overnight culture was grown in LB media from a single transformant. The overnight culture was added to 1L of minimal media optimized for ncAA incorporation^32,86^ and grown at 37 °C until the OD_600_ reached 2.0 and then induced with 0.3 mM isopropyl-b-D-1-thiogalactoside (IPTG). Induction was carried out at 37 °C for 3 hours and the harvested cells were resuspended in 120 ml cell resuspension buffer (30 mM HEPES, pH 7.8, 300 mM KCl, 0.02 % Tween-20, 1.5X protease inhibitor cocktail, 1 mM PMSF, 10 % (v/v) glycerol and 10 mM imidazole). RPA^4AZP^ was purified as described above for unlabeled RPA. To fluorescently label the protein, RPA-DBD-A^4AZP^ or RPA-DBD-D^4AZP^ (∼4 mM in 5 ml storage buffer) was mixed with 1.5-fold molar excess DBCO-Cy5 or DBCO-Cy3 (Click Chemistry Tools Inc.) and incubated for 2 hours at 4 °C. Labeled RPA was resolved from free dye on a Biogel-P4 column (Bio-Rad Inc.) using RPA storage buffer. Fluorescent RPA protein was flash frozen using liquid nitrogen and stored at −70 °C. RPA concentration was measured spectroscopically using ε_280_= 87,210 M^−1^cm^−1^ and labeling efficiency was calculated as described.^86^

### Purification of DSS1

A codon-optimized open reading frame for Human DSS1 (DSS1) was synthesized (GenScript Inc.) with a SUMO protease cleavable N-terminal MVKIH-Strep-6x-HIS-SUMO tag. The MVKIH sequence enhances overall protein overproduction.^87^ The pRSFDuet-1-MVKIH-STREP-SUMO-DSS1 plasmid was transformed into *E. coli* BL21(DE3) pLysS cells and transformants on LB were selected using kanamycin (50 μg/ml). An overnight culture (10 ml) was grown from a single transformant and added to 1 L of LB media containing kanamycin. Cells were grown at 37 °C until the OD_600_ reached 0.6 and then induced with 1 mM IPTG. Induction was carried out at 37 °C for 3 hours and the harvested cells were resuspended in 120 ml cell resuspension buffer (30 mM HEPES, pH 7.8, 300 mM KCl, 0.02 % Tween-20, 1.5X protease inhibitor cocktail, 1 mM PMSF, 10 % (v/v) glycerol, 1 mM TCEP-HCl, and 10 mM imidazole). Cells were lysed using 0.4 mg/ml lysozyme followed by sonication. Clarified lysates were fractionated on a Ni^2+^-NTA agarose column. DSS1 was eluted using cell resuspension buffer containing 50 mM KCl and 400 mM imidazole. Fractions containing DSS1 were pooled and digested for 18 hours at 4 °C with 1:10 molar excess of SUMO protease. The reaction was concentrated using a 30 kDa Amicon spin concentrator and loaded onto a HiLoad Superdex S200 column and fractionated using DSS1 storage buffer (30 mM HEPES, pH 7.8, 300 mM KCl, 0.25 mM EDTA, 0.02 % Tween-20, 1 mM TCEP-HCl, and 10 % (v/v) glycerol). Purified DSS1 protein was flash frozen using liquid nitrogen and stored at −70 °C. DSS1 concentration was measured spectroscopically using ε_280_ = 17,990 M^−1^cm^−1^.

### In vitro kinase assay

For phosphorylation of RPA and RPA variants (RPA^S384A^ and RPA^R382Q^) by Aurora B kinase, recombinant human RPA or RPA-variants (2 μM) were incubated with 250 nM of recombinant Aurora B kinase (Invitrogen Inc.) in kinase reaction buffer (50 mM Tris-HCl, pH 8.0, 10 mM MgCl_2_, 2 mM TCEP, 10 μM ATP and 20 μCi ^32^P-γ-ATP (Perkin Elmer Inc.). Reactions (50 μl total) were initiated by adding kinase and reactions were incubated at 28 ºC for 30 min. 10 μl of 5X Laemmli buffer was added, boiled for a minute, and 30 μl of the reaction was resolved using 8-12 % SDS-PAGE and imaged using Coomassie staining and autoradiography. For analysis of the phosphorylation reaction using mass spectrometry, experiments were performed similarly except 500nM of RPA and variants were used in the reaction along with 50 μM cold-ATP instead of radiolabeled ATP. Reactions were quenched by adding 10 mM EDTA (final concentration) and by freezing on dry ice and analyzed using MS/MS.

### MS/MS analysis of RPA phosphorylation

*In vitro* kinase reactions were subjected to TCA/acetone (10 % TCA in 50 % Acetone final v/v) precipitation for 30 minutes on ice. Precipitated proteins were spun for 10 minutes at room temperature (16,000 g) and the pellets were washed twice with cold acetone. Protein extracts were re-solubilized and denatured in 15 μl of 8 M Urea in 50 mM NH_4_HCO_3_ (pH 8.5) and subsequently diluted to 60 μl for the reduction step with 2.5 μl of 25 mM DTT and 42.5 μl of 25 mM NH_4_HCO_3_ (pH8.5). The diluted reaction was further incubated at 56 °C for 15 minutes and cooled on ice to room temperature. 3 μl of 55 mM chloroacetamide (CAA) was added for alkylation and incubated in darkness at room temperature for 15 minutes. Reaction was quenched by adding 8 μl of 25 mM DTT. Finally, 6 μl of Trypsin/LysC solution (100 ng/μl 1:1 *Trypsin* (Promega) and *LysC* (FujiFilm)) mix in 25 mM NH_4_HCO_3_] and 23 μl of 25 mM NH_4_HCO_3_ (pH8.5) was added to 100 µl final volume. Digestion was carried out for 2 hours at 42°C and an additional 3 µl of trypsin/LysC mix was then added and digested overnight at 37 °C. The reaction was terminated by acidification with 2.5 % trifluoroacetic acid (TFA) to 0.3 % final. Digests were desalted using Agilent Bond Elut OMIX C18 SPE pipette tips per manufacturer protocol, eluted in 10 µl of 70/30/0.1% ACN/H_2_O/TFA, and dried to completion using a speed-vac and finally reconstituted in 25 µl of 0.1 % formic acid. Peptides were analyzed by nanoLC-MS/MS using a Agilent 1100 nanoflow system (Agilent) connected to hybrid linear ion trap-orbitrap mass spectrometer (LTQ-Orbitrap Elite, Thermo Fisher Scientific) equipped with an EASY-Spray electrospray source (held at constant 35 °C). Chromatography of peptides prior to mass spectral analysis was accomplished using a capillary emitter column (PepMap C18, 3 µM, 100 Å, 150×0.075 mm, Thermo Fisher Scientific) onto which 2 µl of extracted peptides was automatically loaded. NanoHPLC system delivered solvents A: 0.1 % (v/v) formic acid, and B: 99.9 % (v/v) acetonitrile, 0.1 % (v/v) formic acid at 0.50 µL/min to load the peptides (over a 30 minute period) and 0.3 µl/min to elute peptides directly into the nano-electrospray with gradual gradient from 0 % (v/v) B to 30 % (v/v) B over 80 minutes and concluded with 5 minute fast gradient from 30 % (v/v) B to 50 % (v/v) B at which time a 5 minute flash-out from 50-95 % (v/v) B took place. As peptides eluted from the HPLC-column/electrospray source survey MS scans were acquired in the Orbitrap with a resolution of 120,000 followed by CID-type MS/MS fragmentation of 30 most intense peptides detected in the MS1 scan from 350 to 1800 m/z; redundancy was limited by dynamic exclusion.

MS/MS data files were converted to mgf file format using MSConvert (ProteoWizard: Open Source Software for Rapid Proteomics Tools Development). Resulting mgf files were used to search against Uniprot *Escherichia coli* proteome databases (UP000000625 01/17/2019 download, 4,446 total entries) containing user defined construct sequences along with a cRAP common lab contaminant database (116 total entries) using in-house Mascot search engine 2.2.07 (Matrix Science) with fixed Cysteine carbamidomethylation and variable Serine and Threonine phosphorylation, Methionine oxidation, plus Asparagine or Glutamine deamidation. Peptide mass tolerance was set at 15 ppm and fragment mass at 0.6 Da. Protein annotations, significance of identification, and spectral based quantification was done with Scaffold software (version 4.11.0, Proteome Software Inc., Portland, OR). Peptide identifications were accepted if they could be established at greater than 96.0 % probability to achieve an FDR less than 1.0 % by the Scaffold Local FDR algorithm. Protein identifications were accepted if they could be established at greater than 99.0 % probability to achieve an FDR less than 1.0% and contained at least 2 identified peptides. Protein probabilities were assigned by the Protein Prophet algorithm.^88^ Proteins that contained similar peptides and could not be differentiated based on MS/MS analysis alone were grouped to satisfy the principles of parsimony. All of the Mascot assigned phophopeptides were subsequently manually interrogated to confirm the identification and residue assignment.

### Cell lines

HCT116 and 293T cells were purchased from American Type Culture Collection. HCT116 cells were maintained in McCoy’s 5A (modified) medium supplemented with 10 % fetal bovine serum (FBS), and 100 units/ml penicillin and 100 mg/ml streptomycin (Gibco). 293T cells were maintained in Dulbecco’s Modified Eagle Medium supplemented with 10 % FBS, 100 units/ml penicillin, and 100 mg/ml streptomycin (Gibco).

### CRISPR/Cas9-mediated gene editing

The RPA-S384A lines were created by the Genome Engineering & Stem Cell Center (GESC@MGI) at Washington University in St. Louis. Briefly, synthetic gRNA targeting the sequence (5’-tccaccgaaatcagagactc<u>NGG</u>) and donor single stranded oligodeoxynucleotides (cttttacaggctgataaatttgatggttctagacagcccgtgttggctatcaaaggagcGcgagtc-Gctgatttcggtggacggagcctctccgtgctgtcttcaagcactatcattgcgaatcc) used for knock-in were purchased from IDT, complexed with Cas9 recombinant protein and transfected into HCT116 cells. Transfected cells were then single cell sorted into 96-well plates, and single cell clones were identified using Next Generation Sequencing to analyze the target site region as those harboring knock-in mutation. Positive clones were expanded, and genotype confirmed prior to cryopreservation. Two clones were selected for this study. All clones were negative for mycoplasma contamination and authenticated as HCT116 cells by STR profiling.

### Western blotting

Cells were washed twice in phosphate buffered saline (1X PBS) and lysed in mammalian cell lysis buffer (MCLB) (50 mM Tris-Cl pH 8.0, 5 mM EDTA, 0.5 % Igepal, 150 mM NaCl) that was supplemented with the following inhibitors just before lysis: 1 mM phenylmethylsufonyl fluoride (PMSF), 1 mM sodium fluoride, 10 mM β-glycerophosphate, 1 mM sodium vanadate, 2 mM DTT, 1X protease inhibitor cocktail (Sigma–Aldrich), 1X phosphatase inhibitor cocktail (Santa Cruz Biotechnology). Lysates were rocked for 15 min at 4°C and cleared by centrifugation at 14,000 rpm for 10 min at 4 °C. Proteins were separated on SDS-PAGE and transferred onto nitrocellulose membranes (0.45 μm; Bio-Rad Laboratories). Membranes were blocked for 1 hr in 5% nonfat dry milk dissolved in Tris Buffered Saline with 0.1% Tween 20 (TBS-T). β-Tubulin (Cell Signaling, 2128), Vinculin (Cell Signaling, 13901), anti-RPA70 (Cell Signaling; 2198), p53 (Santa Cruz Biotechnology, sc126), Histone H3 (1:2000; Cell Signaling, 14269). All phospho-specific primary antibodies were diluted (1:1000) in 1% nonfat dry milk dissolved in TBS-T buffer and rocked overnight at 4°C. The following phospho-specific antibodies were used: pS384-RPA70 (monoclonal, custom generated by Genscript), pS10-Histone H3 (Cell Signaling, 53348), pS139-H2AX (Millipore Sigma, 05-636), pS317-Chk1 (Cell Signaling, 12302). Membranes were probed with HRP-conjugated secondary antibodies (Jackson ImmunoResearch) diluted 1:30000 in TBS-T buffer and incubated for 1 hr at room temperature except for the secondary antibodies targeted against pS384-RPA70 that were diluted in 1% to 2% nonfat dry milk dissolved in TBS-T buffer. Membranes were developed using ECL substrate (Pierce) and chemiluminescence was captured using iBright CL1500 imager (Thermo Fisher).

### Cell viability assays

5,000 HCT116 cells expressing RPA-WT or RPA-S384A were cultured per well of a 96-well plate in McCoy’s culture medium without phenol red. 20 μL of MTS solution (CellTiter 96 Aqueous One solution reagent, Promega) was added to each well, and the plates were incubated for 1 h. Absorbances were read at 490 nm (Synergy H1, BioTek). To obtain background-corrected absorbances, average absorbance values of media with MTS only were subtracted from all other absorbances. To determine rate of cell proliferation, final absorbances were normalized to 0 hrs absorbance of the respective cells. Apoptosis was measured using the Caspase-Glo® 3/7 assay system (Promega). Cells were seeded at a density of 1 × 10^4^ in 96-well black polystyrene microplates (Corning). 100 μL of caspase-Glo reagent, including caspase-Glo substrate, and caspase-Glo buffer was added to each well and incubated for 90 minutes at RT in the dark. Luminescence was measured using a multi-mode microplate reader (Synergy H1, BioTek). To induce replication stress, cells were treated with 10 ng/μL of SN-38 dissolved in DMSO (Tocris). To obtain background-corrected absorbances, average absorbance values of media with reagent only were subtracted from all other absorbances.

### Cell cycle analysis

To synchronize HCT116 cells in mitosis (prometaphase), 1.5 × 10^6^ cells were seeded per 100mm dish. Next day, cells were treated with 75ng/mL nocodazole (Sigma-Aldrich) and incubated for 18 hrs. Rounded mitotic cells were collected by shake-off method and centrifuged at 2000 rpm for 4 min at 4 °C. Cells were washed twice with cold PBS. The cell pellets were lysed in MCLB buffer as described above and collected for western blot analysis. To release cells from mitotic arrest into G1 phase, mitotic cells were collected by shake-off and washed twice in PBS and resuspended in culture media. Cells were then plated onto 100mm dishes and incubated for 3 hrs. Control cells were treated with vehicle-0.1 % DMSO. For inhibition of Aurora kinase B, cells were arrested in mitosis as described above and treated with 3mM Aurora kinase B inhibitor (AZD1152) or 0.1 % DMSO (vehicle) for an additional 45 min. Rounded mitotic cells were collected as described above and centrifuged at 2000 rpm for 2 to 4 min at 4 °C. Cells were washed twice with cold 1XPBS and lysed in MCLB buffer. For cell synchronization using double thymidine block, 1.2 x10^6^ HCT116 cells were seeded per 100 mm dish, and next day, cells were treated with 2 mM thymidine (Sigma-Aldrich) for 16 hrs. Cells were then washed twice with 1XPBS and cultured in media without inhibitors for 8 hrs. For the second thymidine block, cells were cultured in media supplemented with 2 mM thymidine for 16 hrs. Cells were released from G1/S phase arrest by washing twice with 1X PBS and cultured in media without inhibitors. Cells were collected at 3 hrs and 6 hrs post-release from G1/S phase arrest.

### Immunofluorescence

HCT116 cells were grown on poly-d-lysine–coated glass coverslips (Neuvitro) in a 12-well cell culture plate. To arrest cells in mitosis, cells were treated with either 0.1% DMSO (vehicle) or 75 ng/ml nocodazole (Sigma-Aldrich) for 18 hrs. To release cells from mitotic arrest, rounded cells were collected by mitotic shake-off and centrifuged at 2000 rpm for 3 min at RT. Cells were washed twice with PBS with gentle inversion and resuspended in culture media. Cells released from mitotic arrest were grown on poly-d-lysine–coated glass coverslips (Neuvitro) for 1 hr at 37 °C. Cells were fixed in 4 % paraformaldehyde overnight at 4℃. For immunostaining, cells were permeabilized with 2 % Triton X-100 diluted in 1X PBS for 15 min and incubated in blocking buffer (2 % BSA, 0.1 % Igepal, 1X PBS) for 30 min. The following primary antibodies diluted in blocking buffer were used: α-Tubulin (1:45; Cell Signaling, 2125), pS10-Histone H3 (1:1000; Millipore Sigma, 05-1336). Cells were incubated with antibodies in a humidified chamber at RT for 1 hr. Coverslips were rinsed in wash buffer (0.1% Igepal dissolved in 1X PBS) 3 times and the following secondary antibodies diluted in blocking buffer were used: Goat anti-rabbit Alexa Fluor 488 (1:500; Thermo Fisher), Cy3-conjugated AffiniPure donkey anti-mouse (1:200, Jackson ImmunoResearch). Cells were incubated with the secondary antibodies in a humidified chamber at RT for 45 min and washed four times in wash buffer. Coverslips were mounted onto frosted microscope slides (Fisher) using ProLong Gold antifade reagent with DAPI (Invitrogen). Immunostained cells were analyzed under Leica DM6 B upright fluorescent microscope with a 100X oil immersion objective. The digital images were acquired using Leica DFC 9000GT camera and processed by LAS X imaging software. Greater than 70 cells in anaphase/telophase were counted to calculate the frequency of lagging chromosomes and anaphase bridges. For surface area measurements of pS10 Histone H3 stained nuclei, greater than 200 nuclei from three independent replicates were analyzed using ImageJ software (NIH).

### Flow cytometry

Adherent cells were trypsinized and resuspended in culture media. Cells were centrifuged at 1200 rpm for 2 min at 4°C. Mitotic cells were collected by shake-off method as described above. Cell pellets were washed once with cold 1X PBS. Cells were then gently resuspended in 500 μL cold PBS. The cell suspension was added to 5 mL of 100% ethanol dropwise with mild vortexing and stored at -20°C. To stain with propidium iodide (PI), fixed cells were centrifuged at 2000 rpm for 2 min, and pellets were resuspended in 1% bovine serum albumin (BSA) diluted in 1X PBS. Cells were counted and resuspended in PI solution containing 3/50 volume of 50X PI (Sigma-Aldrich), 1/40 volume of 10 mg/mL RNAaseA (Thermo Scientific) and 1% BSA diluted in 1X PBS. Samples were filtered through 35 um strainer caps (Corning) and incubated in dark for 30 min at RT. Samples were analyzed on a BD FACSCanto II flow cytometer using BD FACSDiva software (BD Biosciences). Samples were collected on a low flow rate, and a minimum of 15,000 cycling events were recorded. Data analyses were performed using ModFit LT software (Verity Software).

### Measurement of DNA binding kinetics using stopped-flow fluorescence

All stopped flow experiments were performed on a SX20 instrument (Applied Photophysics Inc.) at 25 °C in RPA reaction buffer (30 mM HEPES pH 7.8, 100 mM KCl, 5 mM MgCl_2_, 6 % (v/v) glycerol, and 1 mM b-ME). The respective mixing schemes are denoted by cartoons alongside the respective figure panels. Seven to eight individual shots were averaged for each experiment. All experiments were repeated a minimum of n=3 times and the mean value and SEM calculated from the individual fits are reported. To capture RPA binding to ssDNA using intrinsic Trp fluorescence, RPA (100 nM) from one syringe was mixed with increasing concentrations of (dT)_35_ ssDNA (0-800 nM) from the other syringe and the change in Trp fluorescence was measured as a function of time by exciting the sample at 290 nm and collecting fluorescence emission using a 305 nm long-pass filter. Data were fit to a single exponential equation and a plot of the k_obs_ (s^−1^) as a function of [RPA] yielded k_on_ and k_off_ values.

A Förster resonance energy transfer (FRET) based experiment was developed to investigate facilitated exchange activity of RPA. Here, assembly of multiple fluorescent RPA molecules on a single ssDNA substrate is captured. RPA molecules are labeled on either DBD-A with Cy5 or DBD-D with Cy3. When these molecules are situated adjacent to each other, an increase in FRET is observed. To measure facilitated exchange, these high-FRET fluorescent RPA containing filaments were challenged with unlabeled RPA and the change in FRET was measured. 200 nM RPA-DBD-A^Cy5^ and 200 nM RPA-DBD-D^Cy3^ were premixed with 90 nM (dT)_97_ and shot against unlabeled RPA (500 nM). Samples were excited at 535 nM (Cy3 wavelength) and Cy5 emission was captured using a 645 nm long-pass filter. Data were fit to s single exponential plus linear equation.

### Hydrogen-Deuterium Exchange mass spectrometry (HDX-MS) analysis

Stock solutions of RPA (13.4 mg/mL) were mixed in the presence or absence of (dT)_35_ ssDNA in a 1:1.2 ratio. Reactions were diluted 1:10 into deuterated reaction buffer (30 mM HEPES, 200 mM KCl, pH 7.8). Control samples were diluted into a non-deuterated reaction buffer. At each time point (0, 0.008, 0.05, 0.5, 3, 30 h), 10 µL of the reaction was removed and quenched by adding 60 µL of 0.75 % formic acid (FA, Sigma) and 0.25 mg/mL porcine pepsin (Sigma) at pH 2.5 on ice. Each sample was digested for 2 min with vortexing every 30 s and flash-frozen in liquid nitrogen. Samples were stored in liquid nitrogen until the LC-MS analysis. LC-MS analysis of RPA was completed as described ^46^. Briefly, the LC-MS analysis of RPA was completed on a 1290 UPLC series chromatography stack (Agilent Technologies) coupled with a 6538 UHD Accurate-Mass QTOF LC/MS mass spectrometer (Agilent Technologies). Peptides were separated on a reverse phase column (Phenomenex Onyx Monolithic C18 column, 100 × 2 mm) at 1 °C using a flow rate of 500 μl/min under the following conditions: 1.0 min, 5% B; 1.0 to 9.0 min, 5 to 45% B; 9.0 to 11.8 min, 45 to 95% B; 11.8 to 12.0 min, 5% B; solvent A = 0.1 % FA (Sigma) in water (Thermo Fisher) and solvent B = 0.1% FA in acetonitrile (Thermo Fisher). Data were acquired at 2 Hz s^−1^ over the scan range 50 to 1700 m/z in the positive mode. Electrospray settings were as follows: the nebulizer set to 3.7 bar, drying gas at 8.0 L/min, drying temperature at 350 °C, and capillary voltage at 3.5 kV. Peptides were identified as previously described^89-92^ using MassHunter Qualitative Analysis, version 6.0 (Agilent Technologies), Peptide Analysis Worksheet (ProteoMetrics LLC), and PeptideShaker, version 1.16.42, paired with SearchGUI, version 3.3.16 (CompOmics). Deuterium uptake was deter- mined and manually confirmed using HDExaminer, version 2.5.1 (Sierra Analytics). Heat maps were created using MSTools.^93^

### RPA-ssDNA binding measured using fluorescence anisotropy

5’-FAM-(dT)_20_ or 5’-FAM-(dT)_40_ were diluted to 10 nM in DNA binding buffer (50 mM Tris acetate pH 7.5, 50 mM KCl, 5 mM MgCl_2_, 1 mM DTT, 10 % (v/v) glycerol, and 0.2 mg/ml BSA). 1.2 ml of this ssDNA stock was taken in 10 mm pathlength quartz cuvettes (Firefly Inc.) maintained at 23 °C and G-factor corrected fluorescence anisotropy was measured (in triplicate) using a PC1 spectrofluorometer (ISS Inc.). Samples were excited at 488 nm and the resulting fluorescence emission was collected using a 520 nm bandpass emission filter. To extract corrected anisotropy from raw values (instrument readings) the concentrations of ssDNA, total fluorescence, and the added protein concentration were first corrected for the effect of sample dilution resulting from the stepwise addition of the protein. Second, the FAM anisotropy was corrected for changes in the fluorescence quantum yield of the bound species due to proximity of the fluorescein moiety to the protein (often referred to as protein-induced fluorescence enhancement or PIFE) in order to plot the anisotropy change due reduction in rotational correlation time alone *i*.*e*., increase in molecular weight of the ssDNA due to complex formation. For this, the dilution-corrected FAM fluorescence values were used to correct and rescale anisotropy values as described.^75^ The saturation points were taken as the intersection of biphasic or triphasic curves from the linear fits of the initial data points reflecting the change in anisotropy upon binding of sub-saturating amounts of proteins.

### Circular Dichroism measurements

CD measurements were performed using a Chirascan V100 spectrometer (Applied Photophysics Inc.). A nitrogen fused set up with a cell path of 1 mm was used to perform the experiments at 20 °C. All CD traces were obtained between 200-260 nm, and traces were background corrected using CD reaction buffer (100 mM NaF, 1 mM TCEP-HCl, and 5 mM Tris-HCl pH 7.5). 600 nM of RPA, RPA^R382Q^, RPA^S384A^ or RPA^S384D^ was used, and 10 scans were collected and averaged per sample using 1 nm step size and 1 nm bandwidth.

### Bio-layer interferometry (BLI) to capture DSS1-RPA interactions

BLI experiments were performed using a single channel BLItz instrument (Sartorius) in advanced kinetics mode at 25 °C with shaking at 2200 rpm. For protein binding, streptavidin (SA) biosensors were pre-hydrated by incubating the tips in BLI buffer (30 mM HEPES pH 7.8, 100 mM KCl, 5 mM MgCl_2_, and 6 % (v/v) glycerol) for 10 min. The experiment was performed as sequential steps in BLI buffer: i) initial baseline was recorded for 30 sec, ii) the tip was incubated with 300 nM of Strep-tagged DSS1 for 120 sec, iii) tip was washed with buffer for 30 sec, iv) binding to different concentrations of RPA was assessed for 120 sec, and v) dissociation was recorded with buffer for 120 sec. Sensorgrams were normalized to the buffer signal. All experiments were repeated at least three times.

### Size exclusion chromatography

SEC experiments to capture formation of RPA nucleoprotein filaments on ssDNA were performed using 600 μl of 3 μM RPA in the absence or presence of ssDNA (3 µM (dT)_35_ or 1 µM (dT)_97_). RPA and ssDNA were premixed and incubated for 10 min on ice and resolved using a Superose 6 Increase (10/300) size exclusion column (Cytiva) equilibrated with SEC buffer (30 mM HEPES, pH 7.8, 100 mM KCl, 1 mM TCEP-HCl, and 10 % (v/v) glycerol). 0.5 ml fractions were collected and assessed using SDS-PAGE.

## Funding

This work was supported by grants from the NIGMS, National Institutes of Health to S.O. (R15 GM126477 and R01 GM143179), and E.A. (R01 GM130746 and R01 GM133967). AUC experiments were supported by a grant from the Office of the Director, National Institutes of Health, S10 OD030343 to E.A. Funding for the Proteomics, Metabolomics and Mass Spectrometry Facility at MSU was made possible in part by the MJ Murdock Charitable Trust and NIGMS of the National Institutes of Health under Award Number P20 GM103474 and S10 OD28650 to B.B. S.K. is supported by a F99/K00 grant from the National Cancer Institute (F99 CA274696).

## Acknowledgements

We thank Dr. Grzegorz Sabat (University of Wisconsin Madison-Mass Spectrometry Core), and GESC@MGI at Washington University in St. Louis for their technical support. We thank Dr. Jaigeeth Deveryshetty for assistance with generating the structural model of RPA.

## Notes

### Competing Interest Statement

The authors have declared no competing interest.

