## Supplementary Figures for "An Aurora B-RPA signaling axis secures chromosome segregation fidelity"

### Supplementary Information

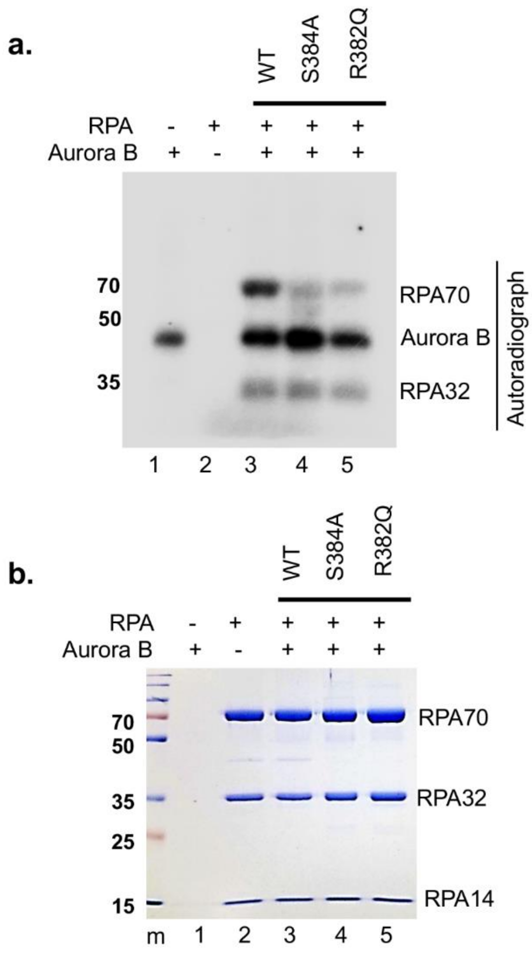


**Supplementary Figure 1. *In vitro* kinase assay shows site-specific phosphorylation of RPA70 on S384 by Aurora B**. **a)** Representative autoradiograph shows Aurora B kinase-dependent phosphorylation of RPA70 that is lost upon phosphosite-Ser384 to Alanine substitution or phospho-motif Arg382 to Glu cancer-specific mutation. Recombinant human RPA was incubated with Aurora B kinase and subjected to *in vitro* kinase assay using $\gamma$-$p^{32}$-ATP. Autophosphorylation of Aurora B kinase is also shown as control. Blots are representative of three independent experiments. **b)** Samples corresponding to **a.** were subjected to SDS-PAGE analysis and Coomassie staining to show equal concentrations of WT-RPA and mutants assayed per reaction. Gels are representative of three independent experiments. m=molecular weight standard.

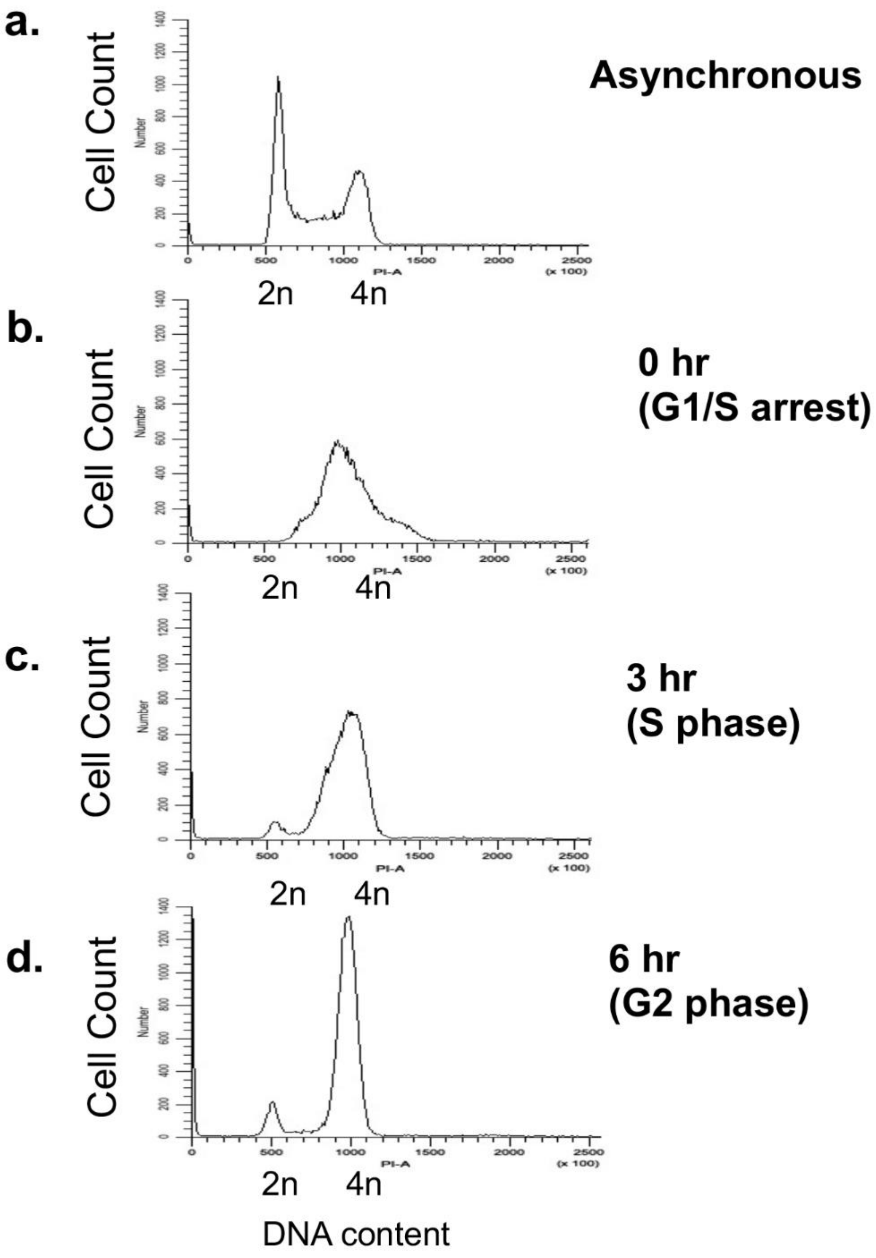


**Supplementary Figure 2. Cell synchronization at G1/S boundary using double thymidine block. a-d)** Cell cycle profile represents synchronization of parental HCT116 cells at the G1/S boundary (**b**) using double thymidine block followed by progression into S phase at 3 hours (**c**) and into G2 phase at 6 hours after release from double-thymidine block (**d**). Asynchronous cells (**a**) used as controls. DNA content was analyzed using propidium iodide staining and flow cytometry. Plots are representative of three independent experiments.

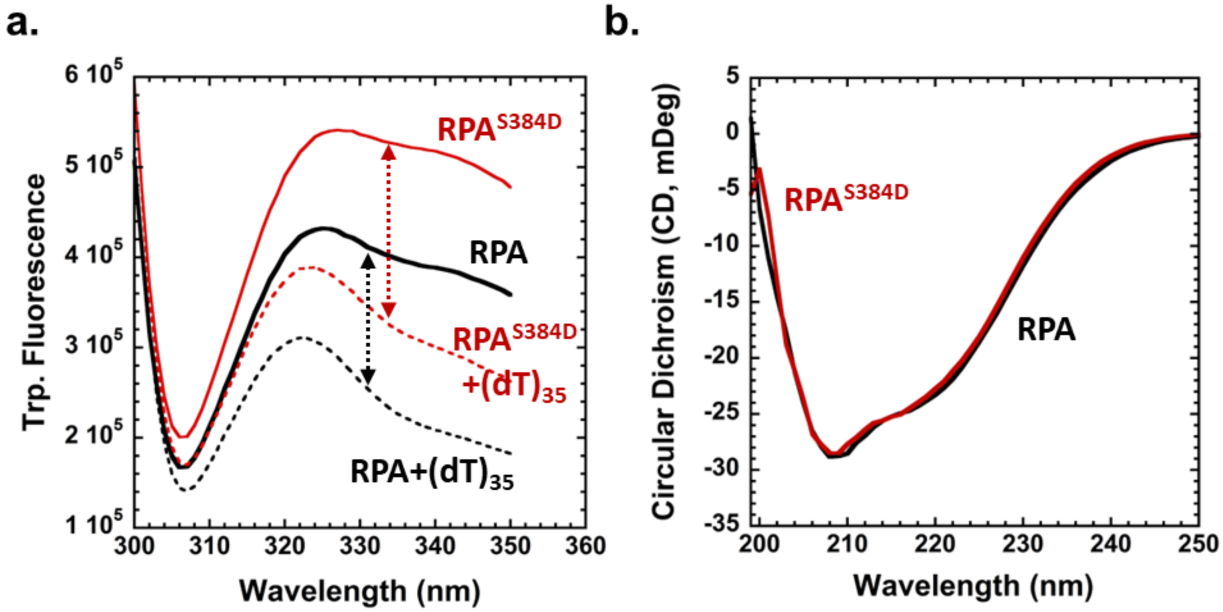


**Supplementary Figure 3. Configuration changes in RPA induced by Aurora B phosphorylation. a)** Changes in intrinsic tryptophan (Trp) fluorescence were collected by exciting RPA or $RPA^{S384D}$ at 295 nm in the presence or absence of $(dT)_{35}$ ssDNA. $RPA^{S384D}$ shows a ~50% enhancement in intrinsic Trp fluorescence compared to RPA. These data show that the domains undergo a configurational change in the phosphomimetic RPA protein leading to a change in the local environment around the Trp residues. Upon ssDNA binding, both RPA and $RPA^{S384D}$ show a similar degree of quenching of Trp fluorescence suggesting no major perturbations in the ssDNA binding properties. **b)** Circular dichroism (CD) analysis of RPA or $RPA^{S384D}$ show no major changes in the overall secondary structures within the domains suggesting that the phosphomimetic substitution does not alter the secondary structure of DBD-D or the other domains in RPA.

a. 3 μM RPA/RPA$^{S384D}$ +/- 3 μM $(dT)_{35}$

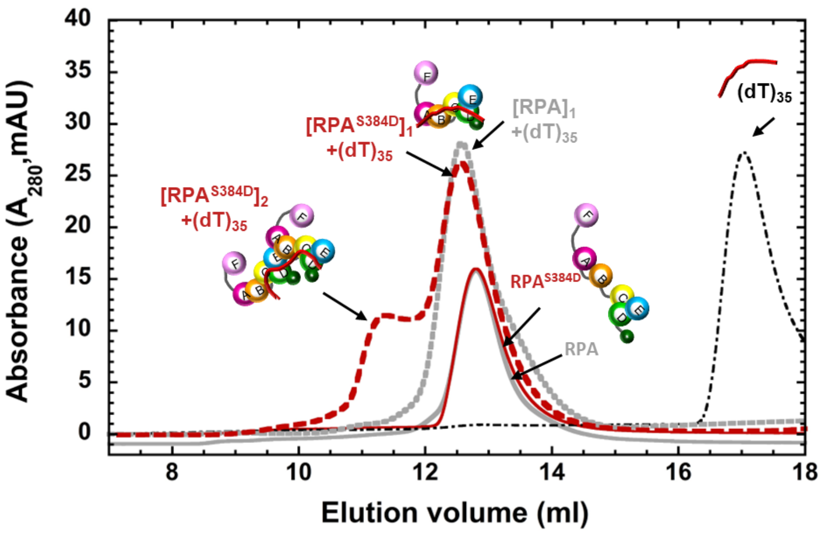


b. 3 μM RPA/RPA$^{S384D}$ +/- 1 μM $(dT)_{97}$

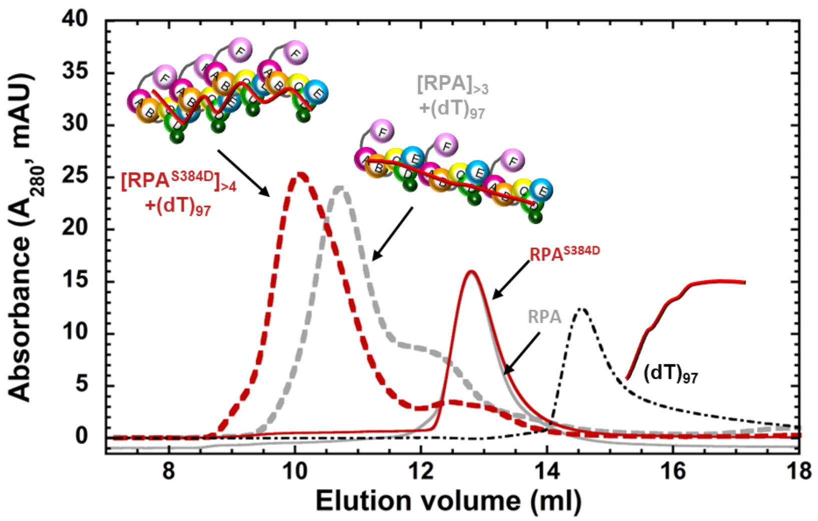


**Supplementary Figure 4. Aurora B phosphorylation promotes formation of higher density RPA-ssDNA nucleoprotein filaments.** RPA binding to **a)** short $(dT)_{35}$ or **b)** long $(dT)_{97}$ ssDNA substrates were assessed by size exclusion chromatography (SEC). RPA and the RPA$^{S384D}$ phosphomimetic elute as single peaks in the absence of ssDNA. On $(dT)_{35}$, incubation of equimolar concentrations of RPA and ssDNA results in a single peak for RPA suggesting formation of a 1:1 complex. However, for RPA$^{S384D}$, a major 1:1 peak is observed along with a minor (larger) 2:1 (RPA$^{S384D}$:DNA) complex peak. This phenomenon is exaggerated on the longer $(dT)_{97}$ substrates where a higher molar ratio of RPA:DNA is used. Here, RPA predominantly forms a 3:1 complex whereas RPA$^{S384D}$ forms a much larger complex which is likely composed of ~4 or 5 RPA$^{S384D}$ bound per $(dT)_{97}$.

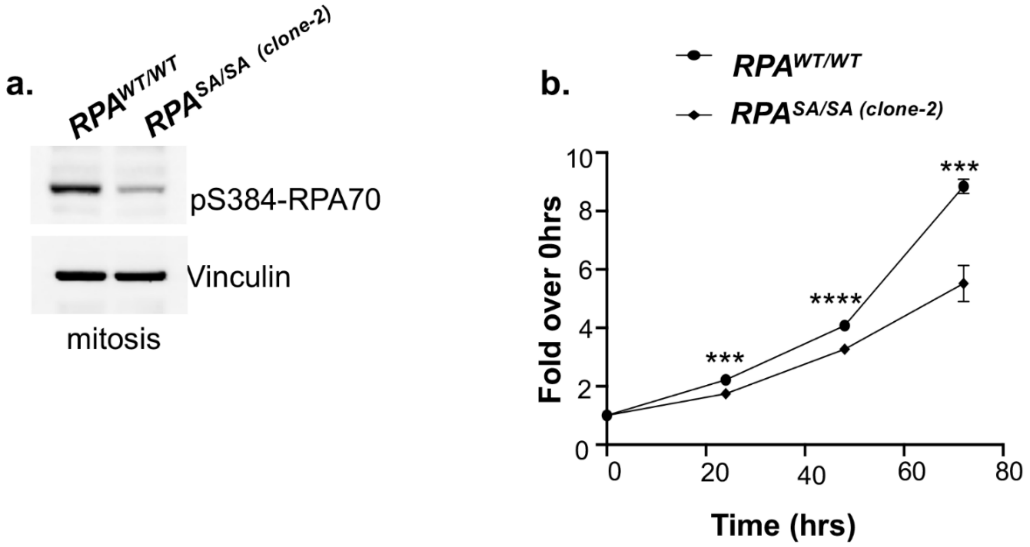


**Supplementary Figure 5. Homozygous knock-in RPA$^{SA/SA}$ mutant clone#2 also exhibits marked loss of viability. a)** Representative western blot depicts loss of ser384-RPA70 phosphorylation in the second clone of RPA$^{SA/SA}$ mutant cells synchronized in mitosis. Blot is representative of three independent experiments. **b)** MTS assay shows decreased viability of phospho-dead *RPA$^{SA/SA}$* mutant. Cells were assayed at 0, 24, 48 and 72 hours of growth. Values corrected for background absorbance were normalized to 0hrs of growth. Error=SEM. Mean of three independent experiments were plotted. Triplicate wells were assayed per time point for each experiment. Statistical significance was determined using an unpaired two-tailed *t*-test: ***$p$=0.0004 at 24 hours, **** $p$<0.0001 at 48 hours, and ***$p$=0.00013 at 72 hours.

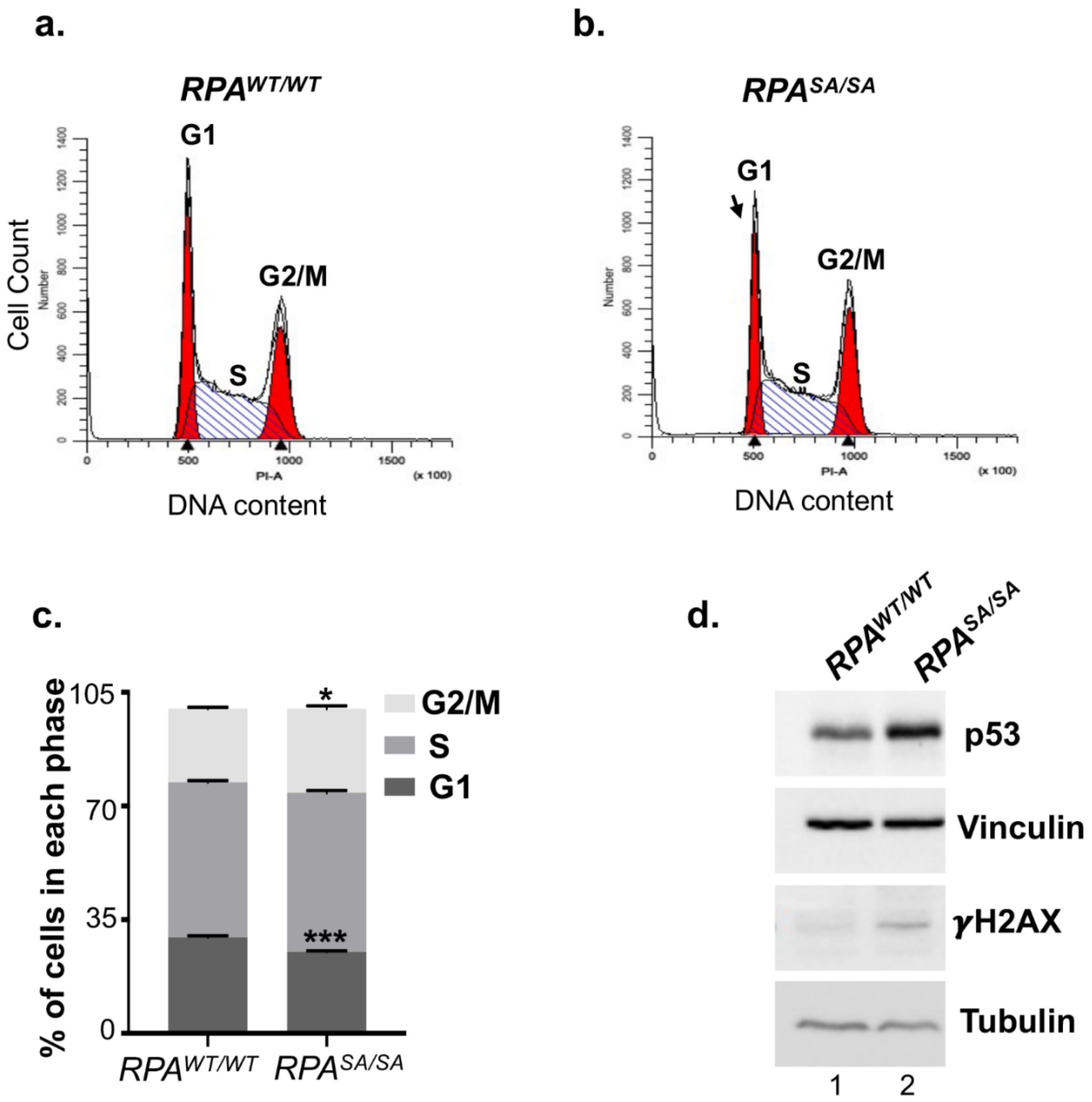


**Supplementary Figure 6. Mild decrease in percentage of cells in G1 phase of cell cycle in the *RPA*$^{SA/SA}$ mutant cells.** Cell cycle profile of asynchronous **a)** *RPA*$^{WT/WT}$ and **b)** *RPA*$^{SA/SA}$ mutant cells were analyzed by flow cytometry and DNA content was assessed by propidium iodide staining. Dead or dying cells were removed by washes and excluded from analysis. Profiles are representative of three independent experiments. **c)** The profiles shown in a. and b. were quantitated and mean cell percentages in each phase of cell cycle were plotted. Bar graph shows mean of three independent experiments. Error = SEM. Statistical significance was determined using an unpaired two-tailed *t*-test: ***$p$=0.0003 (G1), *$p$=0.0245. **d)** Western blot represents basal genomic stress response in *RPA*$^{SA/SA}$ mutant. Blots were probed with the indicated antibodies and are representative of three independent experiments.

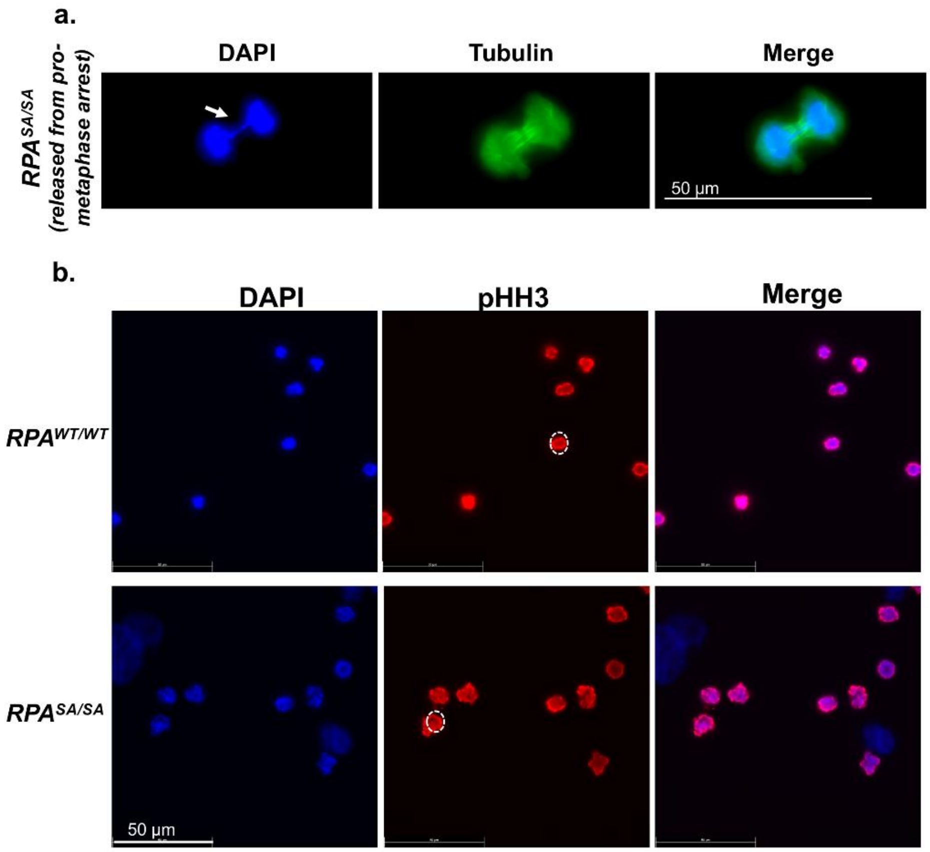


**Supplementary Figure 7. Defects in chromosome segregation and condensation induced by loss of Ser384-RPA70 phosphorylation. a)** Representative immunofluorescent images stained with DAPI, and anti-Tubulin antibody depict anaphase bridges (white arrow) in cells released from prometaphase arrest. Images are representative of three independent experiments. **b)** Representative immunofluorescent images stained with DAPI and anti-phoshpho-Ser10-Histone H3 antibody depict less chromosome condensation (white dotted circles) in cells arrested in prometaphase in *RPA*$^{SA/SA}$ mutant cells.

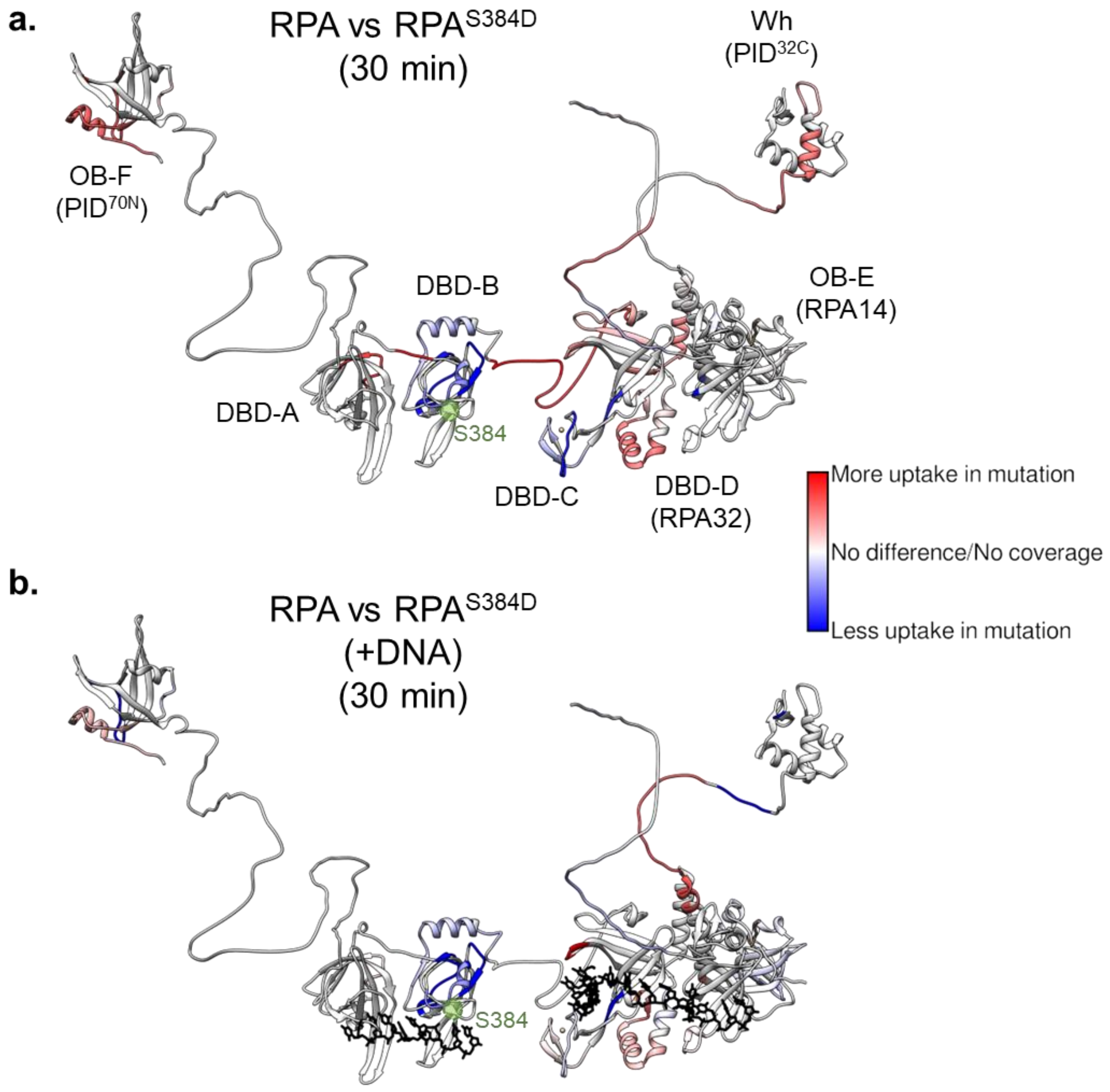


**Supplementary Figure 8. Configurational changes in RPA are induced by a S384D substitution within the Aurora kinase motif in DBD-B.** HDX changes between RPA and $RPA^{S384D}$ are shown in the **a)** absence or **b)** presence of ssDNA. Changes in deuterium uptake/loss are observed in almost all DNA binding and protein-interaction domains. Data are mapped onto the structure of human RPA which is built using the structures of the OB domains from crystal structures. The flexible linkers were modeled using AlphaFold. Position of Ser-384 is denoted in green. Dataset from the 30 min timepoint are shown. Data are averaged from three independent experiments.

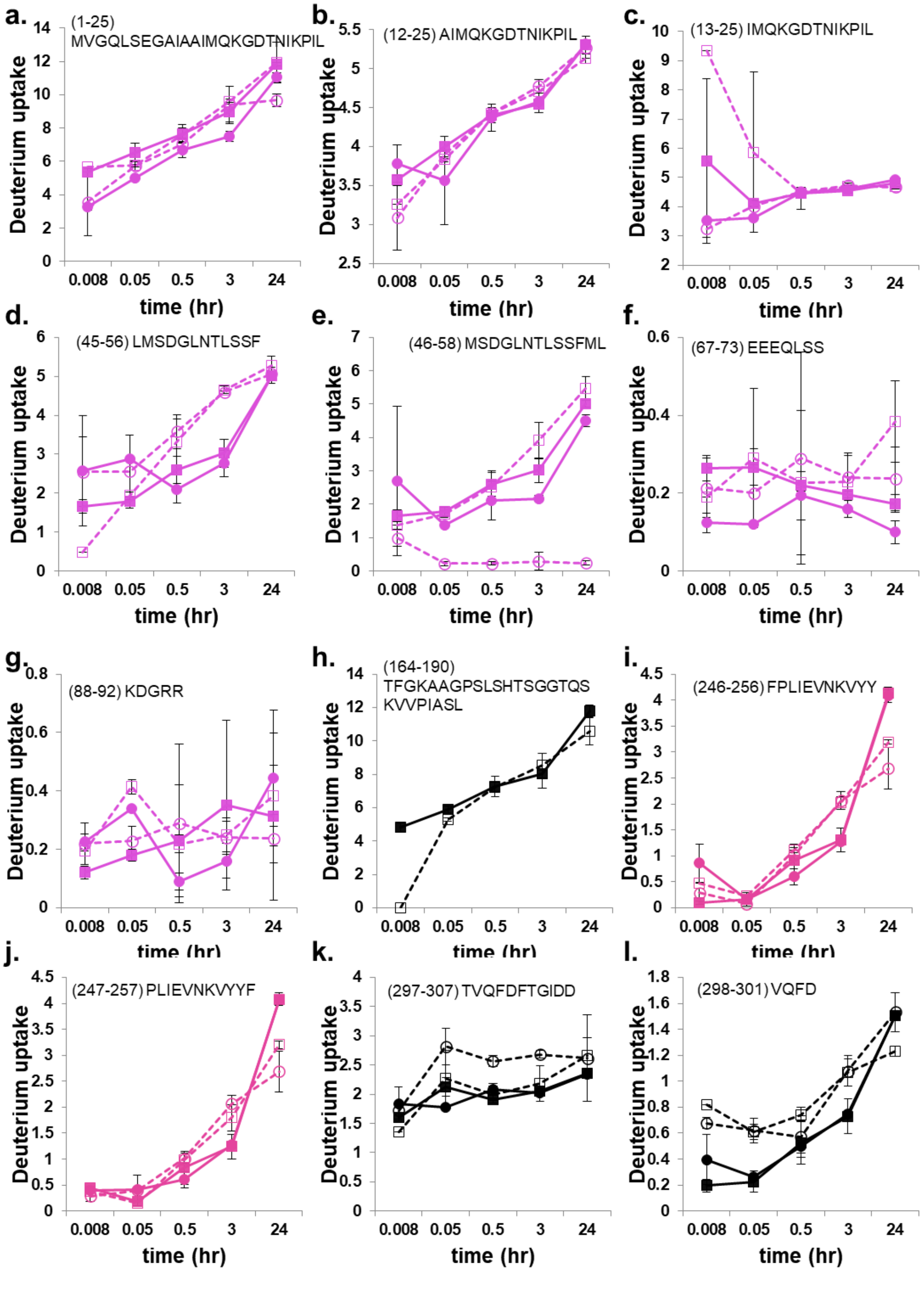


**Supplementary Figure 9. HDX-MS analysis of RPA and RPA$^{S384D}$ peptides from RPA70 in the absence or presence of ssDNA.** HDX-MS data corresponding to specific peptides from RPA and RPA$^{S384D}$ are shown for samples measured in the absence (solid lines) or presence of ssDNA ($(dT)_{35}$)) (dotted lines). Symbols denote RPA (●), RPA+DNA (○), RPA$^{S384D}$ (■), and RPA$^{S384D}$+DNA (□). Peptides from OB-F or PID$^{70N}$ are shown in violet and DBD-A are shown in pink. The data in black are peptides from the F-A and A-B linkers. The amino acid residue numbers and sequence of the corresponding peptides are noted. Std. Dev. from n=3 is plotted.

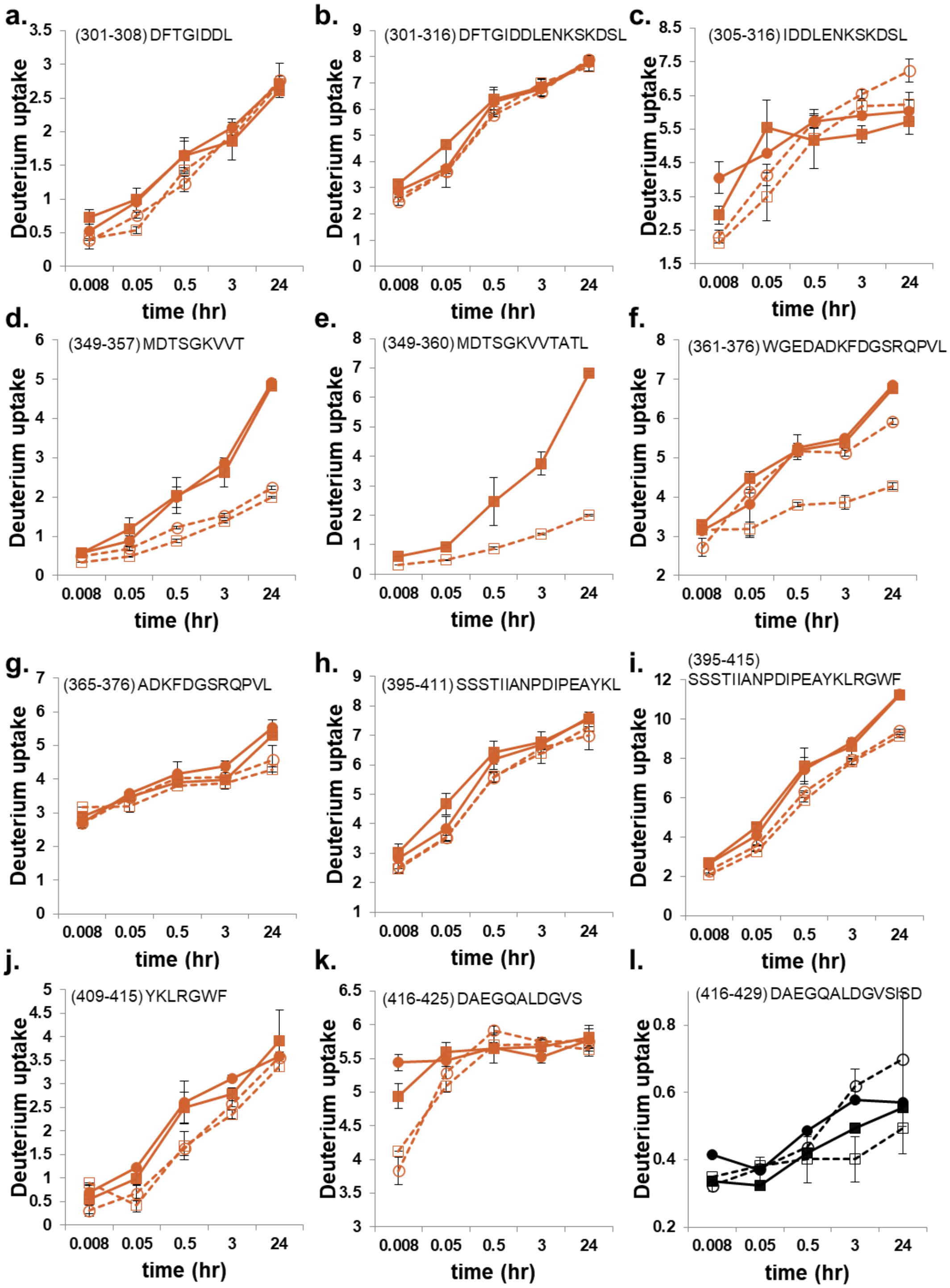


**Supplementary Figure 10. HDX-MS analysis of RPA and RPA$^{S384D}$ peptides from RPA70 in the absence or presence of ssDNA.** HDX-MS data corresponding to specific peptides from RPA and RPA$^{S384D}$ are shown for samples measured in the absence (solid lines) or presence of ssDNA (($(dT)_{35}$)) (dotted lines). Symbols denote RPA (●), RPA+DNA (○), RPA$^{S384D}$ (■), and RPA$^{S384D}$+DNA (□). Peptides from DBD-B are shown in orange. The data in black are peptides from the B-C linker. The amino acid residue numbers and sequence of the corresponding peptides are noted. Std. Dev. from n=3 is plotted.

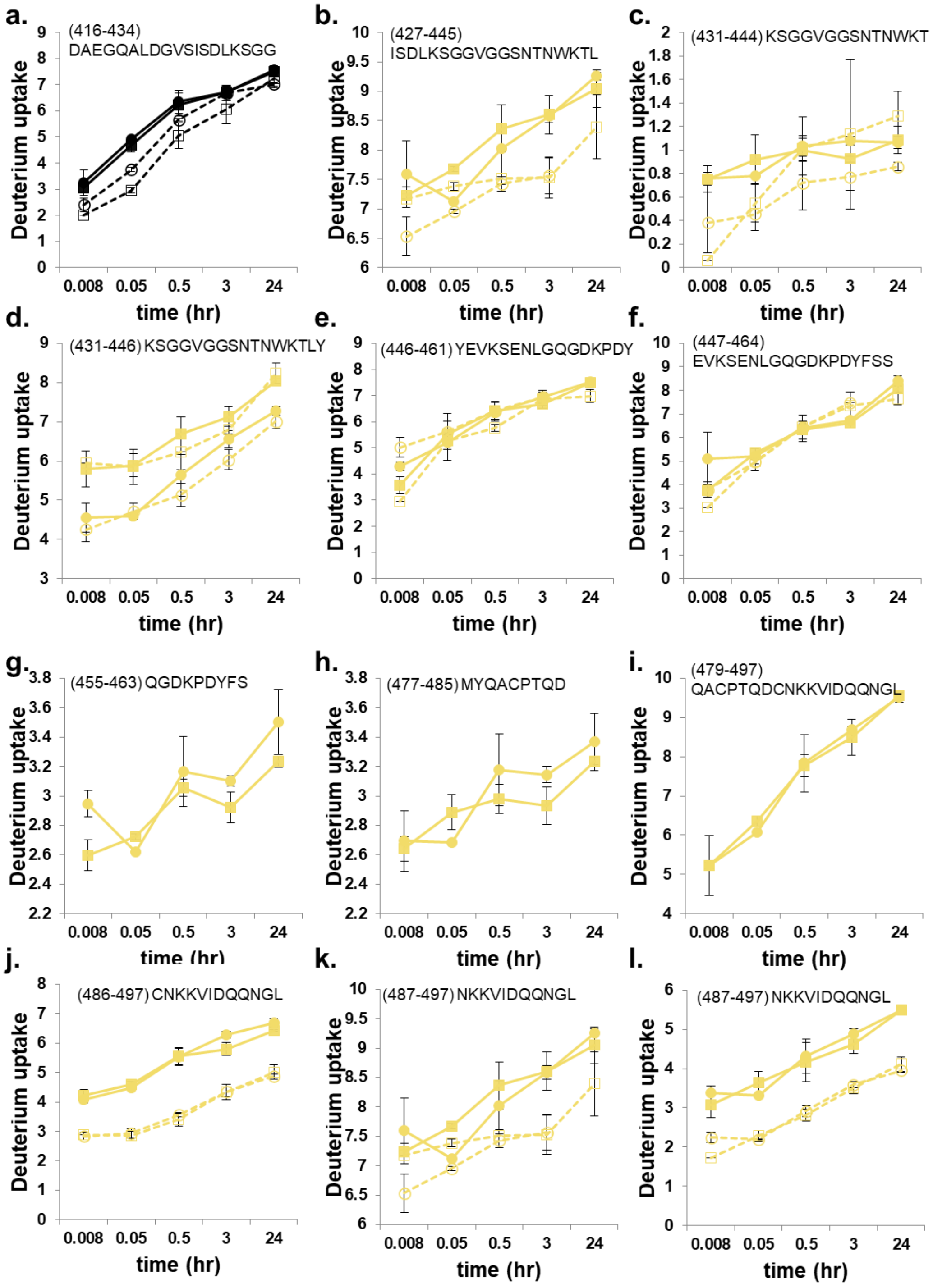


**Supplementary Figure 11. HDX-MS analysis of RPA and RPA$^{S384D}$ peptides from RPA70 in the absence or presence of ssDNA.** HDX-MS data corresponding to specific peptides from RPA and RPA$^{S384D}$ are shown for samples measured in the absence (solid lines) or presence of ssDNA ($(dT)_{35}$) (dotted lines). Symbols denote RPA (●), RPA+DNA (○), RPA$^{S384D}$ (■), and RPA$^{S384D}$+DNA (□). Peptides from DBD-C are shown in yellow. The data in black are peptides from the B-C linker. The amino acid residue numbers and sequence of the corresponding peptides are noted. Std. Dev. from n=3 is plotted.

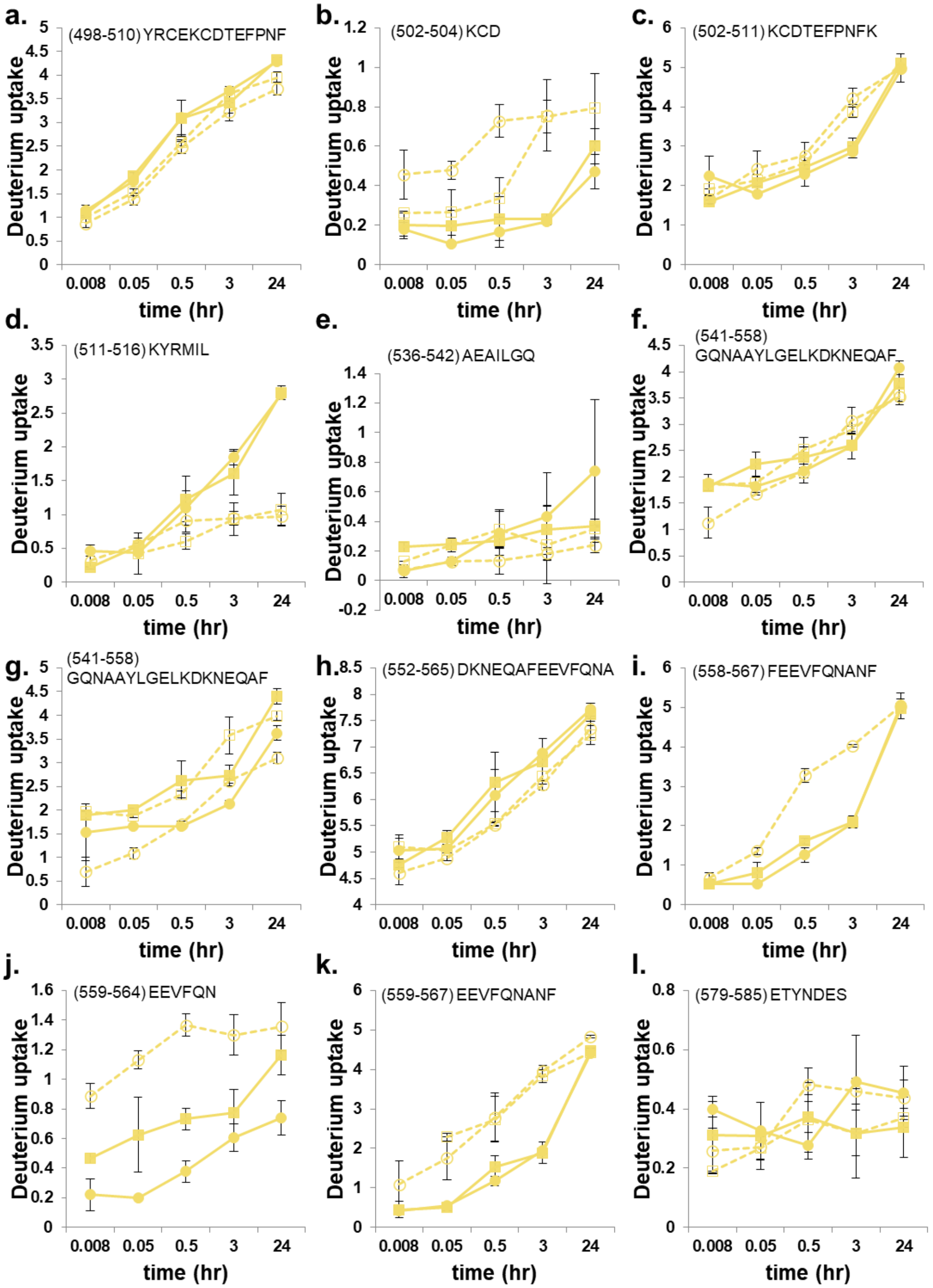


**Supplementary Figure 12. HDX-MS analysis of RPA and RPA$^{S384D}$ peptides from RPA70 in the absence or presence of ssDNA.** HDX-MS data corresponding to specific peptides from RPA and RPA$^{S384D}$ are shown for samples measured in the absence (solid lines) or presence of ssDNA ($(dT)_{35}$) (dotted lines). Symbols denote RPA (●), RPA+DNA (○), RPA$^{S384D}$ (■), and RPA$^{S384D}$+DNA (□). Peptides from DBD-C are shown in yellow. The amino acid residue numbers and sequence of the corresponding peptides are noted. Std. Dev. from n=3 is plotted.

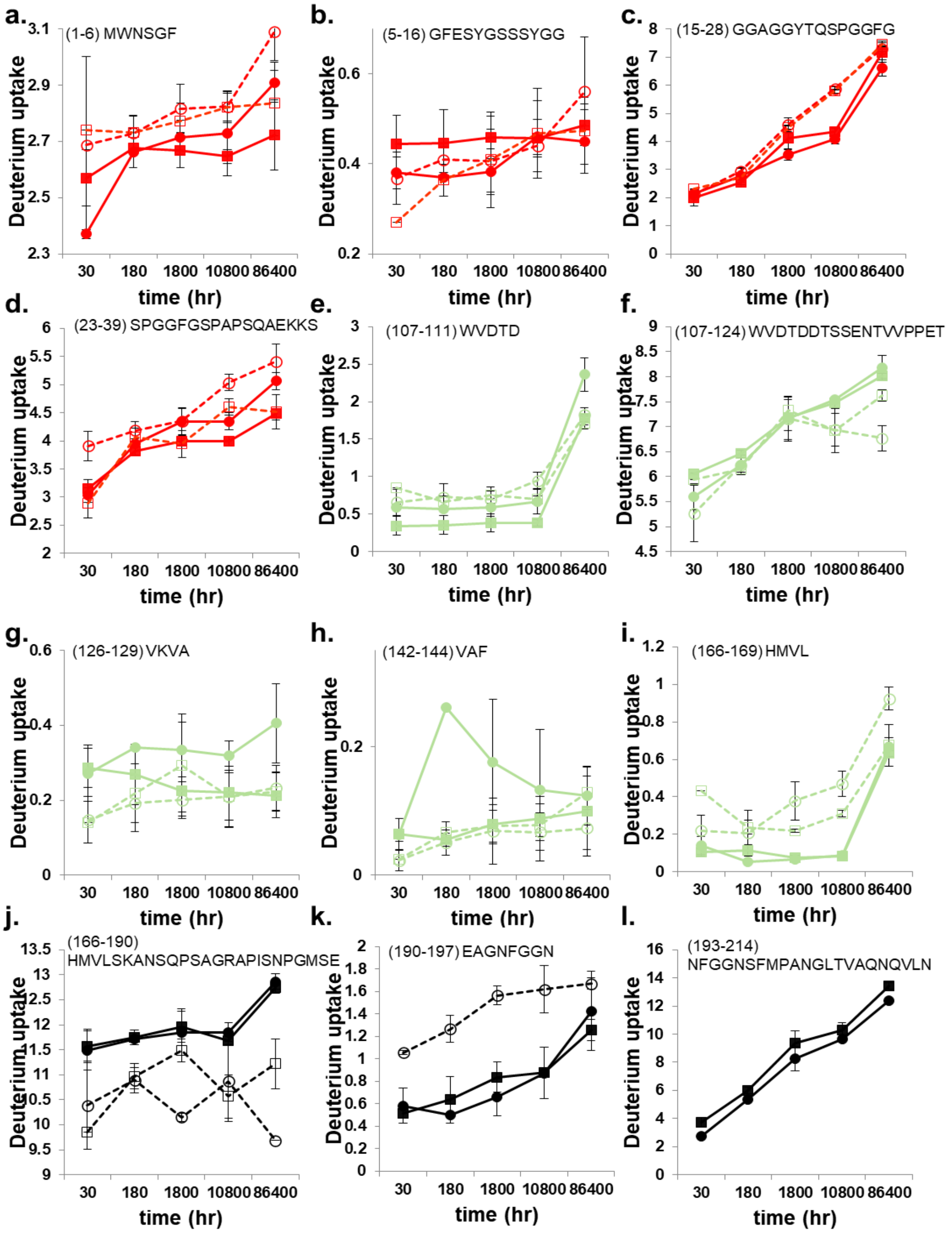


**Supplementary Figure 13. HDX-MS analysis of RPA and RPA$^{S384D}$ peptides from RPA32 in the absence or presence of ssDNA.** HDX-MS data corresponding to specific peptides from RPA and RPA$^{S384D}$ are shown for samples measured in the absence (solid lines) or presence of ssDNA ($(dT)_{35}$) (dotted lines). Symbols denote RPA (●), RPA+DNA (○), RPA$^{S384D}$ (■), and RPA$^{S384D}$+DNA (□). Peptides from the N-terminal hyperphosphorylation region are denoted in red and peptides from DBD-D are shown in green. The data in black are peptides from the D-wh linker. The amino acid residue numbers and sequence of the corresponding peptides are noted. Std. Dev. from n=3 is plotted.

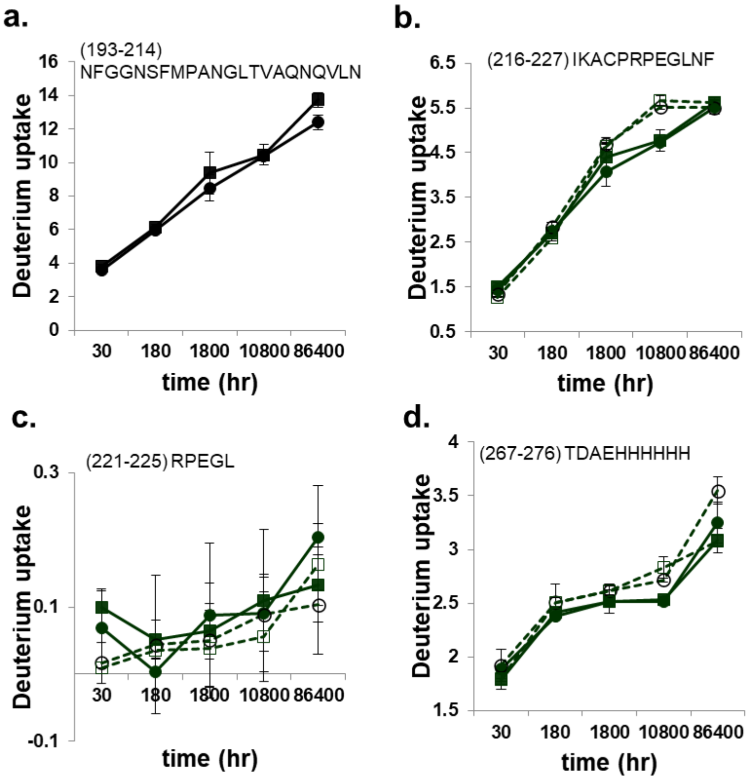


**Supplementary Figure 14. HDX-MS analysis of RPA and RPA$^{S384D}$ peptides from RPA32 in the absence or presence of ssDNA.** HDX-MS data corresponding to specific peptides from RPA and RPA$^{S384D}$ are shown for samples measured in the absence (solid lines) or presence of ssDNA ($((dT)_{35})$) (dotted lines). Symbols denote RPA (●), RPA+DNA (○), RPA$^{S384D}$ (■), and RPA$^{S384D}$+DNA (□). Peptides from the winged helix (wh or PID$^{32C}$) are shown in green. The data in black are peptides from the D-wh linker. The amino acid residue numbers and sequence of the corresponding peptides are noted. Std. Dev. from n=3 is plotted.

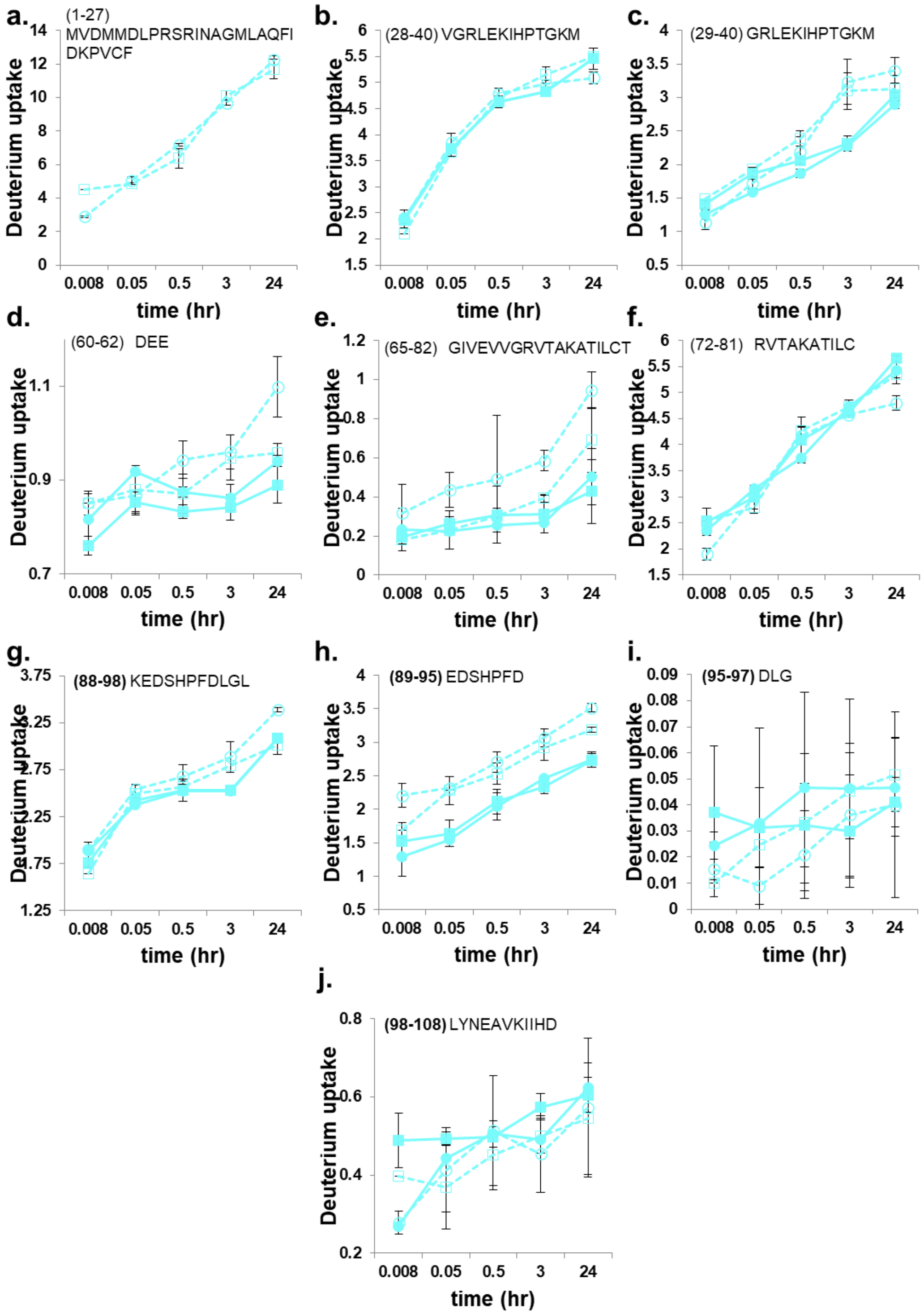


**Supplementary Figure 15. HDX-MS analysis of RPA and RPA$^{S384D}$ peptides from RPA14 in the absence or presence of ssDNA.** HDX-MS data corresponding to specific peptides from RPA and RPA$^{S384D}$ are shown for samples measured in the absence (solid lines) or presence of ssDNA ($(dT)_{35}$) (dotted lines). Symbols denote RPA (●), RPA+DNA (○), RPA$^{S384D}$ (■), and RPA$^{S384D}$+DNA (□). Peptides from RPA14 are shown in cyan. The amino acid residue numbers and sequence of the corresponding peptides are noted. Std. Dev. from n=3 is plotted.
